# Repertoire-wide phylogenetic models of B cell molecular evolution reveal evolutionary signatures of aging and vaccination

**DOI:** 10.1101/558825

**Authors:** Kenneth B Hoehn, Jason A. Vander Heiden, Julian Q. Zhou, Gerton Lunter, Oliver G. Pybus, Steven H. Kleinstein

## Abstract

In order to produce effective antibodies, B cells undergo rapid somatic hypermutation (SHM) and selection for binding affinity to antigen via a process called affinity maturation. The similarities between this process and evolution by natural selection have led many groups to use evolutionary and phylogenetic methods to characterize the development of immunological memory, vaccination, and other processes that depend on affinity maturation. However, these applications are limited by several features of affinity maturation that violate assumptions in standard phylogenetic models. Further, most phylogenetic models are designed to be applied to individual lineages comprising genetically diverse sequences, while B cell repertoires often consist of hundreds to thousands of separate low-diversity lineages. Here, we introduce a hierarchical phylogenetic framework that incorporates the unique features of SHM, and integrates information from all lineages in a repertoire to more precisely estimate model parameters. We demonstrate the power of this approach by characterizing previously undescribed phenomena in affinity maturation. First, we find evidence consistent with age related changes in SHM hot-and cold-spot motifs. Second, we identify a consistent relationship between increased tree length and signs of increased negative selection, apparent in the repertoires of both healthy subjects and those undergoing active immune responses. This suggests that B cell lineages shift towards negative selection over time as a general feature of affinity maturation. Our study provides a framework for undertaking repertoire-wide phylogenetic testing of SHM hypotheses, and provides a new means of charactering the process of mutation and selection during affinity maturation.

## Introduction

B cell receptors (BCRs) are membrane-bound immunoglobulins (Ig) expressed on the surfaces of B cells that bind to antigen and may be released as antibodies to fight infection. BCRs are generated through the shuffling of Ig gene segments by V(D)J recombination and, if the cell expressing them is activated, by a second process of BCR modification called affinity maturation (Murphy *et al.* 2012). Affinity maturation consists of repeated rounds of somatic hypermutation (SHM) of the BCR, cell proliferation, and selection for antigen binding affinity (Murphy *et al.* 2012). These processes give rise to clonal lineages of B cells that each descend from a progenitor cell, from which they differ predominately by point mutations. The BCR sequence, and the nature of the mutations introduced during affinity maturation, can be investigated in detail using high-throughput next generation BCR sequencing (Boyd *et al.* 2009; Greiff *et al.* 2015b; Yaari and Kleinstein 2015).

The fact that affinity maturation is analogous to the process of evolution by natural selection suggests that methods from molecular evolutionary biology, particularly phylogenetics, could have broad utility in studying affinity maturation. This has stimulated the development of methods of evolutionary sequence analysis designed specifically for BCR sequences, particularly phylogenetic approaches (Barak *et al.* 2008; Kepler 2013; Hoehn *et al.* 2017; DeWitt *et al.* 2018). These methods have shown promise in elucidating information about the adaptive immune response in humans, such as the sequence of mutations that occur during antibody co-evolution with HIV (Liao *et al.* 2013) and the migration of B cells in multiple sclerosis (Stern *et al.* 2014).

There are a number of challenges in adapting phylogenetic techniques to B cell clonal lineage analysis. For example, the biology of affinity maturation violates fundamental assumptions in most phylogenetic substitution models, such as independent change at each nucleotide site and time-reversibility of substitution rates (Hoehn *et al.* 2016). To address these issues we previously introduced the HLP17 substitution model, which weights codon substitutions by the presence of SHM hot/cold-spot motifs, and does not assume that substitution rates are time reversible (Hoehn *et al.* 2017).

Phylogenetic techniques are typically used to analyze individual, genetically-diverse B cell lineages (Felsenstein 1981; Liao *et al.* 2013). However, B cell repertoires are typically profiled by next-generation sequencing and consist of many – potentially thousands of – expanded clonal lineages, each of which may contain only a few unique sequences (Weinstein *et al.* 2009; Jiang *et al.* 2013; Greiff *et al.* 2015a). In light of this, many analyses have used non-phylogenetic summary statistics to characterize BCR repertoires, such as the distribution of sequences per clone (e.g. Bashford-Rogers *et al.* 2013; Greiff *et al.* 2015a; Hoehn *et al.* 2015). Others have used non-phylogenetic approaches to study somatic hypermutation biases (Yaari *et al.* 2013) and signatures of clonal selection (Yaari *et al.* 2012), which represent B cell clonal lineages by selecting or constructing a single representative sequence. SHM biases have also recently been incorporated into a phylogenetic framework under Feng *et al.* (2017). By explicitly modeling shared ancestry among sequences within the same clone, phylogenetic approaches potentially offer a more powerful means of understanding somatic hypermutation and affinity maturation by using the full set of substitutions expected to have occurred in a repertoire. However, standard phylogenetic approaches are limited to single lineages, and give imprecise parameter estimates, except when applied to unusually long-lived or highly-diverse B cell lineages (e.g. Hoehn *et al.* 2017).

We propose here that it is possible to combine some of the benefits of phylogenetic and summary statistic approaches of B cell repertoire analysis, by using hierarchical phylogenetic models. These approaches contain multiple levels of parameters, some of which are shared among lineages, while others are estimated for each lineage individually (Suchard *et al.* 2003). For example, Rodrigo et al. (2003) applied one such hierarchical phylogenetic approach to a set of HIV sequences from infected patients in order to jointly estimate both the virus substitution rate and the proportion of individuals that did not respond to antiretroviral therapy. However, previous applications of hierarchical phylogenetic models to virus genomes do not address the abovementioned model assumptions that are violated by the biology of B cell affinity maturation.

A hierarchical approach that is specifically tailored to B cell sequence evolution has the potential to dramatically improve accuracy of parameter estimation. By assuming that B cell lineages within a particular repertoire experience broadly similar patterns of substitution (e.g., hot-and cold-spot sequence motifs that experience altered mutation rates under SHM), a hierarchical approach is able to share information across B cell lineages and thereby take advantage of the large genetic diversity of the entire repertoire, despite the fact that each individual lineage within the BCR repertoire data may exhibit low diversity. Here, we develop a hierarchical phylogenetic framework capable of characterizing entire B cell repertoires by jointly estimating parameters and lineage tree topologies for all lineages within a repertoire. We first introduce the HLP19 model, an improved form of the HLP17 model that accounts for changing codon frequencies during affinity maturation, and validate our hierarchical approach through simulation. We then apply this new framework to characterize the effects of aging on B cell repertoire development and B cell responses to influenza vaccination. We demonstrate that hierarchical approaches can quantify variation in SHM features both across individuals and within the same individual through time. Our results reveal previously uncharacterized immunological phenomena underlying aging and vaccination. We discover (i) evidence of changes in SHM hot/cold-spot mutation biases associated with age, (ii) evidence of negative selection acting on complementarity-determining regions (CDRs) associated with the human immune response to influenza vaccination, and (iii) a consistent relationship between increased lineage tree length and signatures of negative selection across our datasets.

## Methods

### A nonstationary, nonreversible phylogenetic substitution model for B cell evolution

The process of nucleotide change along a given phylogenetic tree is modeled as a Markov process, such that the rate of transitioning into any state at each instant in time is dependent only on the current state of the model (Felsenstein 1981). Here, we characterize codon change in Ig sequences using a 61×61 element matrix (**Q** matrix) that describes the instantaneous rates of change between all non-stop codons. In the HLP17 model (Hoehn *et al.* 2017), these instantaneous rates are parameterized by the nonsynonymous/synonymous mutation rate ratio (ω), transition/transversion mutation rate ratio (κ), codon frequencies (**π**), and modified substitution rates *h*^*a*^ where *a* is an SHM hot-or cold-spot motif, such as WR<u>C</u> (Peled *et al.* 2008); W=A/T, R=A/G; only the underlined base experiences increased substitution).

The HLP17 model makes a salient approximation (as do almost all other substitution models) that nucleotide or codon frequencies are constant over time at a stationary distribution. However, the codon composition of B cell sequences begins substantially far away from equilibrium and changes over time (Sheng *et al.* 2016), making this assumption inappropriate. Hoehn *et al.* (2017) attempted to address this problem by using maximum likelihood (ML) to estimate codon frequencies. Whilst this approach may be better than empirical estimates of codon frequencies, at least in some instances, it more than doubles the number of model parameters. Here, we introduce HLP19, a modified HLP17 substitution model which, among other minor adjustments (see **Supplemental File 1**), uses the predicted codon frequencies at the midpoint of phylogeny in question. HLP19 is fully detailed in **Supplemental File 1**. Overall, HLP19 has less than half the number of free parameters as HLP17 and exhibits improved branch length estimates, generally better estimates of certain substitution model parameters such as ω (**Supplemental File 3**), and is structurally more similar to other non-reversible substitution models (Yang 1994; Kaehler *et al.* 2017). Further, certain approximations in HLP19 (detailed in **Supplemental File 1**) give significantly improved runtime compared to HLP17.

### Hierarchical phylogenetic models for B cell repertoires

Under standard ML phylogenetics (Felsenstein 1981), a single multiple sequence alignment ***X*** is specified, and the goal is to find the tree topology and branch lengths ***T***, and the set of substitution parameters, that maximize the likelihood of ***X***. For B cell lineage phylogenies, the sequence alignment is supplemented with a predicted germline sequence ***G*** that acts as an outgroup and adds direction to the tree. In this study we extend this approach by calculating the likelihood of the entire B cell repertoire, which we define as the product of the tree likelihoods for each of *n* lineages, using each lineage *i*’s tree topology (*T*_*i*_), substitution parameters (ω _*i*_, κ _*i*_, ***h***_*i*_), sequence data (*X*_*i*_) and predicted germline sequence (*G*_*i*_) (equation 1). This approach therefore assumes that mutations in each lineage are independent from each other:

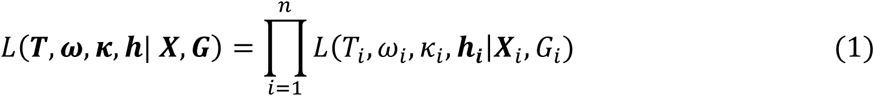

The goal of our phylogenetic repertoire analysis is to find the tree topologies, branch lengths, and substitution parameters that maximize the product on the left-hand side of equation 1, whereas the goal of typical ML phylogenetic analysis is to maximize individually each phylogenetic likelihood on the right-hand side. In hierarchical models, parameters may be constrained to be identical across lineages, allowing them to be estimated at the repertoire level. For instance, we may estimate a repertoire-wide transition/transversion rate ratio by constraining κ_1_ = κ_2_ = … κ_n_. Constraining parameters in this way will lower the overall likelihood of the repertoire compared to optimizing parameters for each lineage individually (because there will be fewer degrees of freedom) but will decrease the number of parameters and thereby reduce parameter estimation variance. For the analyses presented here, we constrain all substitution parameters to be identical across lineages within a repertoire.

### B cell repertoire datasets

We use hierarchical phylogenetic models to characterize B cell repertoires in two previously published data sets obtained from peripheral blood samples. The first dataset (Age) consists of samples taken from 27 healthy individuals in two consecutive years (Wang *et al.* 2013). Subjects varied in age from 20 to 81 years old and both male and female subjects were included. Our second dataset (Vaccine) consists of samples from three male donors at 10 time points: −8 days, −2 days, −1 hour, +1 hour, +1 day, +3 days, +7 days, +14 days, and +28 days relative to seasonal influenza vaccination (Laserson *et al.* 2014). Quality control and data processing for both of these datasets is detailed in **Supplemental File 1**. Samples in the Vaccine dataset had between 141 and 15,763 (mean 6,272.7) unique sequences in non-singleton clones (i.e. clones containing >1 unique sequence). Because of this large variation, the repertoires in the Vaccine dataset were subsampled to a depth of 3,000 sequences in non-singleton clones (**Supplemental File 1**). Sequence depth in the Age dataset was more even, with between 370 and 2,065 (mean 1,126) unique sequences in non-singleton clones, so the repertoires in the Age dataset were not subsampled.

### Phylogenetic model parameter and topology estimation

We used a single-linkage hierarchical clustering approach, detailed in **Supplemental File 1**, to assign sequences into clonal lineages, each of which were assumed to descend from a single naïve B cell ancestor. Because we were not able to reliably predict the junction regions of germline sequences (Gupta *et al.* 2015), we removed the CDR3 from all sequences analyzed. We then used the hierarchical phylogenetic model described above to quantify effects of BCR mutation and selection in the Age and Vaccine datasets.

Phylogenetic model parameters are an important source of information about evolutionary dynamics. For example, the amino acid replacement vs. silent mutation rate ratio (ω) can be used to distinguish positive and negative selection (Nielsen and Yang 1998), while the relative rate of transitions to transversions (κ) can be informative about mutation biases. We first estimated maximum likelihood tree topologies and branch lengths for each B cell lineage using the GY94 (Goldman and Yang 1994; Nielsen and Yang 1998) substitution model, in which single, shared ω and κ parameters were estimated for each repertoire, and codon frequencies were set to their empirical frequencies across all sequences within each repertoire. For computational efficiency, we used these estimated topologies to estimate branch lengths and substitution parameters of the HLP19 model at the repertoire level; namely, we estimated κ, ω_FWR_, ω_CDR_ (separate ω values for CDR and FWRs), and individual *h*^*a*^ values (altered relative mutation rate) for WR<u>C</u>, <u>G</u>YW, W<u>A</u>, <u>T</u>W, SY<u>C</u>, and <u>G</u>RS hot-and cold-spot motifs (see Hoehn *et al.* (2017) for more details on these parameters).

Hypotheses concerning substitution model parameter estimates can be tested in a phylogenetic framework using a likelihood ratio test (Huelsenbeck and Rannala 1997). For models that differ only by one free parameter, a *p* value of 0.05 corresponds to a log-likelihood difference of 1.92 between the alternative (ML estimated) and null (fixed value) model (Huelsenbeck and Rannala 1997). The log-likelihood ratio test allows estimation of 95% confidence intervals for parameter estimates using profile likelihood curves. Each point on a profile likelihood curve is created by calculating the maximum likelihood obtained when the parameter of interest is fixed to a particular value and all other parameters are optimized. We used a straightforward binary search and linear interpolation approach to estimate the 95% confidence interval either side of the ML estimate.

### Dataset simulation

As a means of validation, simulations (detailed in **Supplemental File 2**) were performed to test (i) the performance of the HLP19 model relative to the previous HLP17 and GY94 models (**Supplemental File 3**) and (ii) the effects of estimating parameters using a hierarchical phylogenetic model compared to inference from individual lineage trees (**Supplemental File 4**). To verify that the trends we observe in the Age and Vaccine datasets are not simply the result of biases in our parameter estimation procedure, we performed simulations using pre-specified substitution parameters (κ = 2, ω_FWR_ = 0.4, ω_CDR_ = 0.7, *h*^*WRC*^ = 4, *h*^*GYW*^ = 6, *h*^*WA*^ = 4, *h*^*TW*^ = 2, *h*^*SYC*^ = −0.6, and *h*^*GR*S^ = −0.6) and the same tree topologies and branch lengths as the empirical trees from the Age and Vaccine datasets. We then repeated the analyses performed in each section on these simulated datasets (**Supplemental File 6**).

## Results

### Repertoire-wide phylogenetic models improve parameter estimation

Phylogenetic substitution model parameters can be an important source of information about the evolutionary dynamics of lineages; for instance, the amino acid replacement vs. silent mutation rate ratio (ω) is used to characterize natural selection operating on genetic sequences (Nielsen and Yang 1998). The HLP19 model parameters are informative about the process of B cell affinity maturation. The model includes separate ω parameters for the framework (FWR) and complementarity determining (CDR) regions (ωFWR and ω_CDR_), the transition/transversion rate ratio (κ), and a set of altered substitution rates at SHM hot/cold-spot motifs (*h*^*WRC*^, *h*^*GYW*^, *h*^*WA*^, *h*^*TW*^, *h*^*SYC*^, *h*^*GR*S^; nucleotides represented using the International Union of Pure and Applied Chemistry (IUPAC) coding scheme, only underlined bases experience altered rates).

The small size of most B cell lineages poses a problem for accurate estimation of phylogenetic model parameters for individual B cell lineages. Namely, the size distributions of clonal lineages within B cell repertoires, particularly those derived from blood samples, typically follow a power-law distribution and are dominated by many lineages that each carry only a few unique sequences (Weinstein *et al.* 2009; Jiang *et al.* 2013). We confirmed this pattern using blood sample-derived BCR repertoires from 27 healthy subjects (Age dataset; Wang *et al.* 2013). Across these subjects 88 to 96% (mean: 92.3%) of lineages comprised a single unique sequence, and between 98 to 99.8% (mean: 99.3%) of lineages contained < 5 unique sequences. To test our ability to estimate phylogenetic substitution model parameters from small lineages such as these, we simulated B cell repertoire datasets using a model of SHM sequence evolution, and re-estimated HLP19 model parameters for each lineage individually (**Supplemental File 4**). These results confirm that model parameter estimation is highly inaccurate for most individual B cell lineages within this dataset, with mean proportional error rates ranging from 3.37 for *h*^*<u>G</u>YW*^ to 211.35 for ω_CDR_ (**Figure 1**; **Supplemental File 4**).

**Figure 1:**
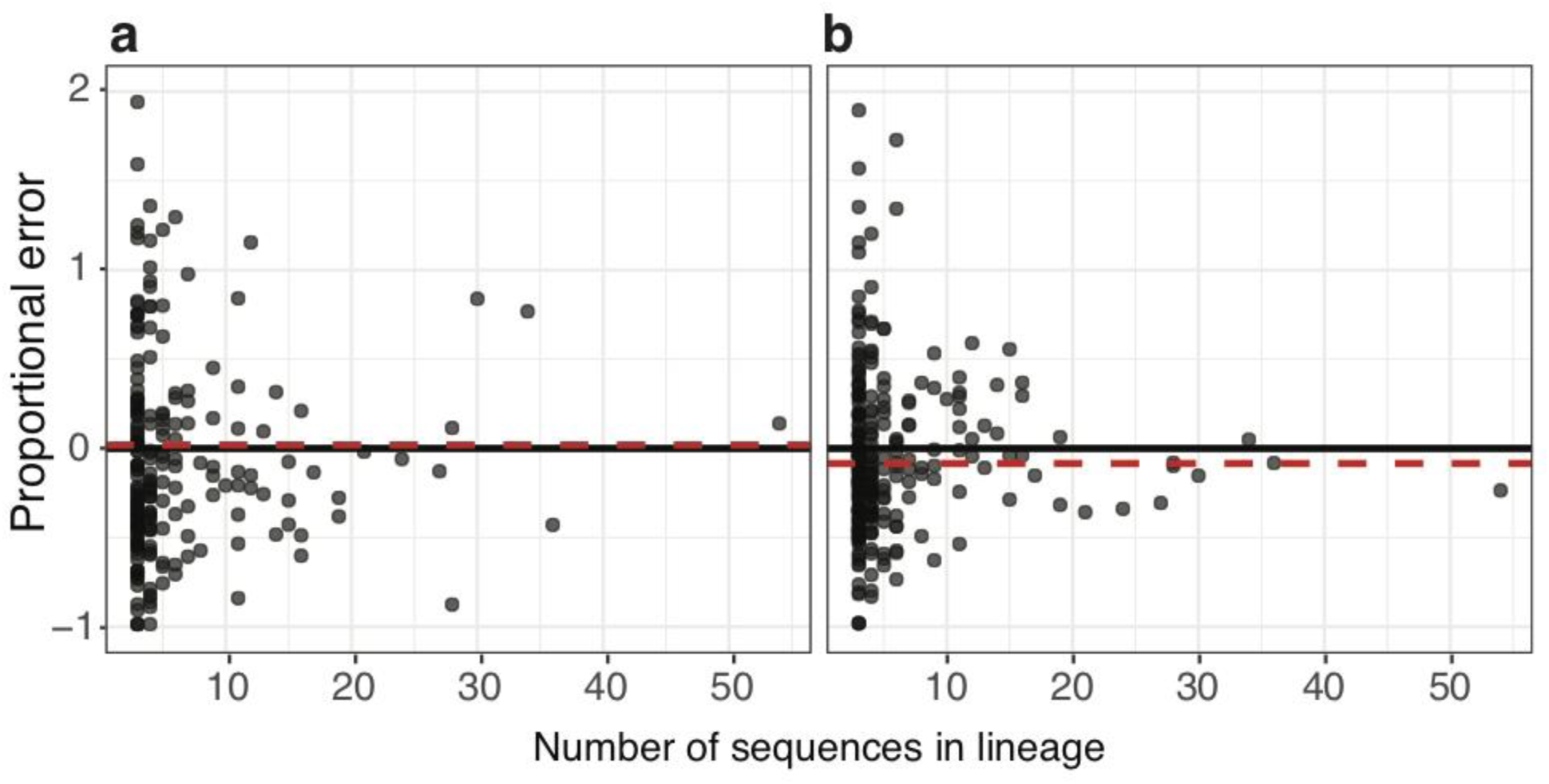
Increased precision of repertoire-wide parameter estimates (**a**) Proportional error in estimates of the ω_CDR_ parameter under the HLP19 model. (**b**) Proportional error in estimates of the ω_FWR_ parameter under the HLP19 model. In both panels, the black dots show the values estimated from each individual lineage B cell lineage and the red lines shows the estimate obtained from all lineages combined using a hierarchical model. Data were generated from a simulated repertoire using tree topologies from subject 97 in the Age dataset (see **Supplemental File 4** for full details and results).

Given the overrepresentation of small lineages in B cell repertoires, we sought to increase the precision of parameter estimation using a hierarchical phylogenetic framework. As outlined in **Methods**, this is done by constraining model parameters to be identical among lineages, and then estimating the set of parameter values that maximizes the likelihood of the entire repertoire.

Applying this hierarchical framework to the same simulation data as above (**Supplemental File 4**), we found that sharing parameters across lineages substantially reduced error in estimates for all substitution model parameters compared to individual estimates, often by multiple orders of magnitude (**Figure 1; Supplemental File 4**). Thus, a repertoire-wide phylogenetic approach leverages the information available across all lineages in the repertoire and more precisely estimates biological parameters relating to affinity maturation.

To test the strengths and weaknesses of different phylogenetic models in our hierarchical framework, we compared the performance of three codon substitution models: GY94 (Goldman and Yang 1994), which does not include SHM hot-or cold-spot motifs, HLP17 (Hoehn et al 2017), which is a modification of the GY94 model that incorporates hot-and cold-spot biases, and HLP19, which is a modification of the HLP17 introduced herein (**Supplemental File 1**) that incorporates codon frequencies in a manner that is more interpretable and formally similar to previous nonreversible models (Kaehler *et al.* 2017). Simulation analyses performed using multiple tree topologies and parameter values (**Supplemental File 3)** revealed that parameter estimates under HLP19 had a lower mean absolute bias across all parameters (0.03) than HLP17 (0.08) and GY94 (0.16; **Supplemental File 3**). For parameters relating to mutation (κ, *h* values), HLP19 showed a slightly lower mean absolute bias (0.04) compared to HLP17 (0.06; **Supplemental File 3**). HLP19 and GY94 models had similar absolute bias in branch lengths (< 0.002), which was lower than that of HLP17 (0.11; **Supplemental File 3**). HLP17 performed worse than GY94 in branch length estimation, which is surprising given Hoehn *et al.* (2017) showed that the HLP17 model improved branch length estimates compared to GY94. However, we have since determined that the simulations performed in Hoehn *et al.* (2017) were unintentionally but unfairly biased towards the HLP17 model (see detailed explanation in **Supplemental File 2)**. The simulations performed here do not have this issue, and show that HLP19 largely addresses the weaknesses of on HLP17 in branch length estimation (**Supplemental File 3**). Perhaps most importantly, for parameters relating to selection (ω_FWR_ and ω_CDR_) HLP19 showed significantly lower mean absolute bias (0.02) compared to HLP17 (0.1) and GY94 (0.29; **Supplemental File 3**). Mean bias of ω_CDR_ estimates were especially high under the GY94 model (range: 0.38 to 0.59) and increased in simulations with higher hot-spot mutation rates and longer branch lengths (**Supplemental File 3**). This echoes previous findings; models that fail to account for altered mutation rates of SHM motifs (e.g. GY94) can significantly bias estimates of ω (dN/dS) in BCR lineages towards detecting positive selection in the CDRs (Dunn-Walters and Spencer 1998; Hershberg *et al.* 2008). Overall, we found the HLP19 model improves upon the GY94 and HLP17 models, particularly when estimating ω_CDR_ and branch lengths, respectively.

To further compare the appropriateness of the GY94, HLP17, and HLP19 models when applied to BCR repertoire data, we estimated how well each model fit our empirical datasets using the Akaike information criterion (AIC; Akaike 1974). The AIC uses the maximum log-likelihood estimated using a model, penalized by the number of freely estimated parameters. Smaller AIC values are generally interpreted as better model fit. In all 27 subjects AIC was highest under GY94 and lowest under HLP19, indicating that the HLP19 model had a significantly better fit to all subjects compared to the GY94 and HLP17 models (**Supplemental File 5**).

### Variation of model parameters within and among subjects

We tested whether repertoire-wide parameter estimates can reproduce known features of somatic hypermutation targeting, such as hot/cold-spot targeting (Yaari *et al.* 2013) by estimating HLP19 model parameters from BCR repertoire data that was obtained from 27 healthy individuals of varying age and sex (Age dataset; Wang *et al.* 2013). While the values of parameter estimates varied, all subjects exhibited the same overall pattern in model parameters that relate to SHM targeting (**Figure 2**). In all subjects, <u>G</u>YW motifs exhibited the largest substitution rate increases of the all motifs considered (*h* ^*GYW*^ values were 4 to 6), followed by the WR<u>C</u> (*h*^*<u>W</u>RC*^ ∼3), W<u>A</u> (*h*^*W<u>A</u>*^ ∼3) and <u>T</u>W (*h*^*<u>T</u>W*^∼1) motifs. Symmetrical SY<u>C</u> and <u>G</u>RS motifs were estimated to be mutational cold-spots (*h*^*SY<u>C</u>*^ and *h*^*<u>G</u>RS*^ ∼ −0.6). We compared these parameter estimates to mutability estimates under the S5F model (Yaari *et al.* 2013), which describes the relative mutation rate of sequence pentamers during SHM in an independent and separate cohort of healthy subjects. When averaging over pentamers within particular SHM motifs under uniform pentamer frequencies, the S5F model predicts the same ranking as we obtained using the HLP19 model: <u>G</u>YW (mean mutability= 2.46) > <u>W</u>RC (1.87) ≈ W<u>A</u> (1.71) > <u>T</u>W (1.19) > SY<u>C</u> (0.23) ≈ <u>G</u>RS (0.22).The transition/transversion rate ratio (κ) estimated by our repertoire-wide model was ∼2, which is also consistent with previous findings (Betz *et al.* 1993; Cowell and Kepler 2000). Overall, these results show that repertoire-wide parameter estimates obtained using a hierarchical phylogenetic approach are broadly consistent with previous expectations in healthy individuals.

**Figure 2:**
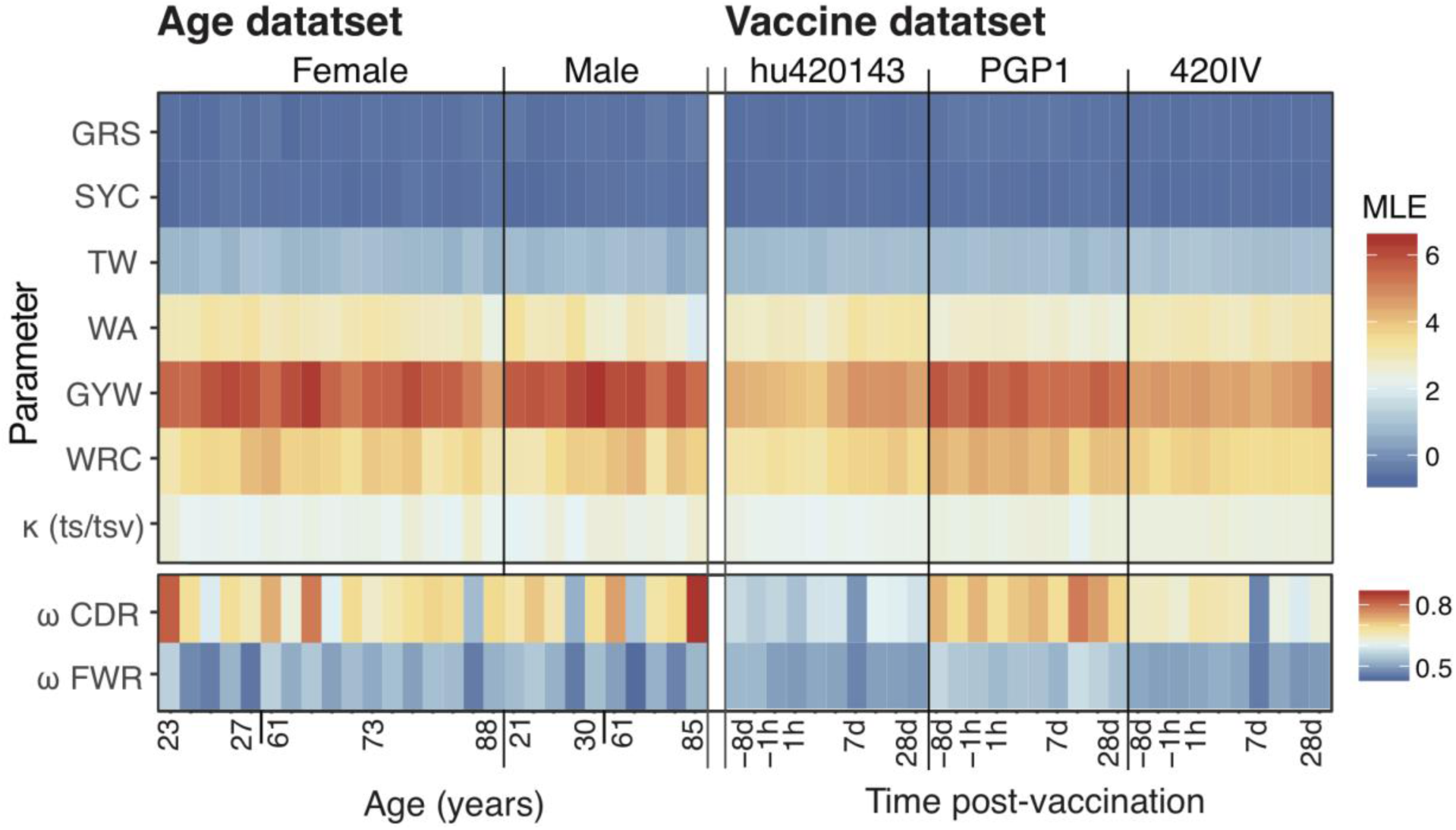
Variation of parameter estimates by subject and time in the Age and Vaccine datasets (**a)** HLP19 parameter estimates from each subject in the Age dataset, ordered by sex and age. (**b**) HLP19 parameter estimates for the Vaccine dataset, ordered by subject and sample time relative to influenza vaccination. The upper box in both (a) and (b) shows the model parameters that relate to somatic hypermutation (motif targeting, transition/transversion ratio). The lower box shows estimates of ω_CDR_ and ω_FWR_, which relate to selection. Note that values in the lower box are scaled differently from those in the upper panel (see legends on right hand side).

### Age is associated with changes in SHM mutation biases

Age and sex are associated with substantial differences in the immune system; for example, older individuals are more vulnerable to infection (Castle 2000; Fink *et al.* 2018), while females are at a higher risk of developing auto-immune diseases (Ansar Ahmed *et al.* 1985). We sought to investigate whether the mutation and selection processes underlying SHM might contribute to these differences.

To investigate potential age-and sex-related differences in SHM targeting, we analyzed the 27 subjects surveyed by Wang et al (2013; Age dataset), which included both male and female subjects with an age range of 21 to 88 years at the time of sampling. We used multiple linear regression to investigate the effects of age and sex on estimated model parameters. Age and sex were modeled as interaction variables against the estimated substitution rate biases of SHM motifs (i.e. the HLP19 model *h* values; **Supplemental File 1**). Because we conducted 18 tests in all (two dependent and nine independent variables), we used Benjamini-Hochberg (Benjamini and Hochberg 1995) multiple hypothesis test correction to adjust *p*-values. Substitution rates in W<u>A</u> (*h*^*W<u>A</u>*^) were significantly negatively associated with age in both male (coefficient=-0.011; adjusted p=0.001), and female subjects (coefficient=-0.006; adjusted p=0.03). No other parameter showed a significant interaction with age in either sex after *p*-value adjustment. These results are consistent with a model in which older individuals have reduced mutation bias towards W<u>A</u> hot-spots, possibly reflecting a difference in SHM mechanism in these individuals.

We performed simulation analyses to test whether the observed trends between *h*^*WA*^ and age could be due to biases in our parameter estimation procedure (*Methods*; **Supplemental File 6**). For all 20 simulated repetitions of the Age dataset, the *h*^*WA*^ slope coefficients for males and females were closer to zero than their respective empirical estimates (**Supplemental File 6a**). These results demonstrate that these trends are due to factors other than biases in parameter estimation, given the underlying structure of our datasets and predicted germline sequences.

### Variation in signatures of selection is uncorrelated with age, sex, EBV, and CMV status

Antigen-driven selection plays a major role in shaping BCR repertoire diversity. In molecular evolutionary biology, selective dynamics are often characterized by estimating the relative rate of substitutions that change amino acids versus those that do not, often called dN/dS or ω (Nielsen and Yang 1998). Low ω values are indicative of fewer amino acid changes than expected, which is generally interpreted as resulting from negative selection. We estimate ω separately for the complementarity determining regions (CDRs) and framework regions (FWR). Estimates of ω_FWR_ are expected to be lower than those of ω_CDR_ because FWRs are more structurally constrained than CDRs (Shlomchik *et al.* 1989), which are primarily used in antigen binding (Murphy *et al.* 2012; Yaari *et al.* 2012). Consistent with this expectation, we found that in the Age dataset, estimated ω_CDR_ values (range: 0.52 – 0.87, mean: 0.68) were higher than estimated ω_FWR_ values (range: 0.44 – 0.56, mean: 0.51) in all 27 subjects (*p* < 0.001; paired Wilcoxon test; **Figure 2**). ω_CDR_ estimates were also more varied among subjects than ω_FWR_ values, perhaps representing different individual histories of antigenic stimulation. However, we were unable to find a clear biological correlate of ω_CDR_ in the Age dataset among the variables provided with the data (Wang *et al.* 2013). Specifically, values of ω_CDR_ did not show a significant relationship with age (slope *p*-value = 0.66; least squares regression; **Figure 3**), sex (*p* = 1.0; Wilcoxon rank sum test), Epstein-Barr virus seropositivity (*p* = 0.19; Wilcoxon rank sum test), or cytomegalovirus seropositivity (*p* = 0.19; Wilcoxon rank sum test).

**Figure 3:**
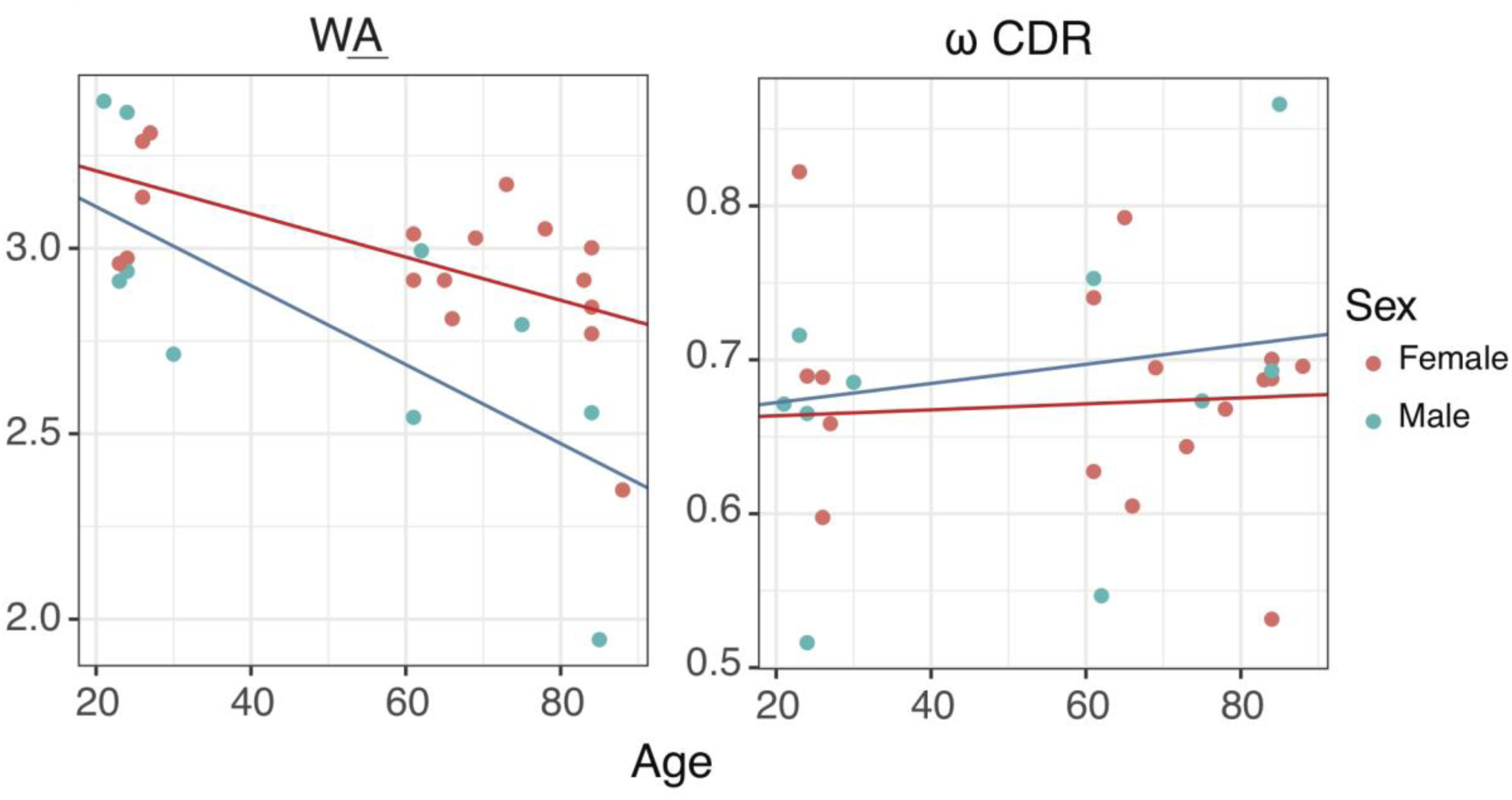
Change in SHM motif mutation rates with age. Linear regressions of age against model parameters under the HLP19 model, estimated using maximum likelihood. Separate regressions are shown for male (blue) and female (red) subjects. The *h*^*WA*^ parameter shows a significant decrease with age in males and females.

### Post-influenza vaccination repertoires show signs of negative selection and longer tree length

Influenza vaccination substantially perturbs the B cell repertoire. A large, antigen-specific plasmablast response is observed in the blood ∼7 days post-vaccination which subsides approximately one week later (Wrammert *et al.* 2008; Jackson *et al.* 2014). To investigate the selective dynamics of this process, we estimated HLP19 substitution model parameters using the repertoires of three otherwise healthy subjects who were sampled 10 times over the course of influenza vaccination, beginning 8 days prior to vaccination and ending 28 days afterwards (Laserson *et al.* 2014). Because we were primarily interested in selection and genetic diversity of these samples, we focused on changes in ω_CDR_, the relative rate of nonsynonymous/synonymous substitutions, and tree length (the total expected substitutions per site within an individual lineage phylogeny).

We found a variety of responses among the subjects. PGP1, the oldest subject of the three, did not show any clear patterns of change over time, in either mean tree length or ω_CDR_. Notably, this subject at day +14 had only 141 sequences and consequently very wide 95% confidence intervals, illustrating the importance of correctly estimating model uncertainty in analysis of BCR sequence data.

In contrast to PGP1, subjects 420IV and hu420143 both showed increased tree length at day+7 compared to one hour prior to vaccination (-1h), consistent with the expected burst of BCR genetic diversity 7 days post-vaccination (Wrammert *et al.* 2008). The estimated mean tree length within a sample was highest at day+7 for subjects 420IV and hu420143, with a fold increase of 2.38 and 1.18 compared to one hour prior to vaccination (-1h) (**Figure 4**). Consistent with this, multiple large clones in subjects 420IV and hu420143 arose at day+7 (**Supplemental File 8**). In addition to increased tree length, day+7 was associated with a significant decrease in ω_CDR_ in both these subjects (**Figure 4**). For 420IV at -1h, ω_CDR_ = 0.64 (95% CI: 0.6, 0.66) and at day+7 ω_CDR_ = 0.47 (95% CI: 0.45, 0.50). For hu420143 at -1h, ω_CDR_ = 0.57 (95% CI: 0.54, 0.59) and at day +7 ω_CDR_ = 0.49 (95% CI: 0.46, 0.51). Interestingly, though 420IV and hu420139 had different pre-vaccination estimates of ω_CDR_ (0.64 and 0.57, respectively) their estimates were similar at day+7 (0.47 and 0.49), day+14 (0.62 and 0.61), and day+21 (0.60 and 0.60; **Figure 4**). Overall, this indicates that, at the expected date of peak vaccine response, the repertoires of these two subjects were characterized by an increase in genetically-diverse BCR lineages and signatures of increased negative selection.

**Figure 4:**
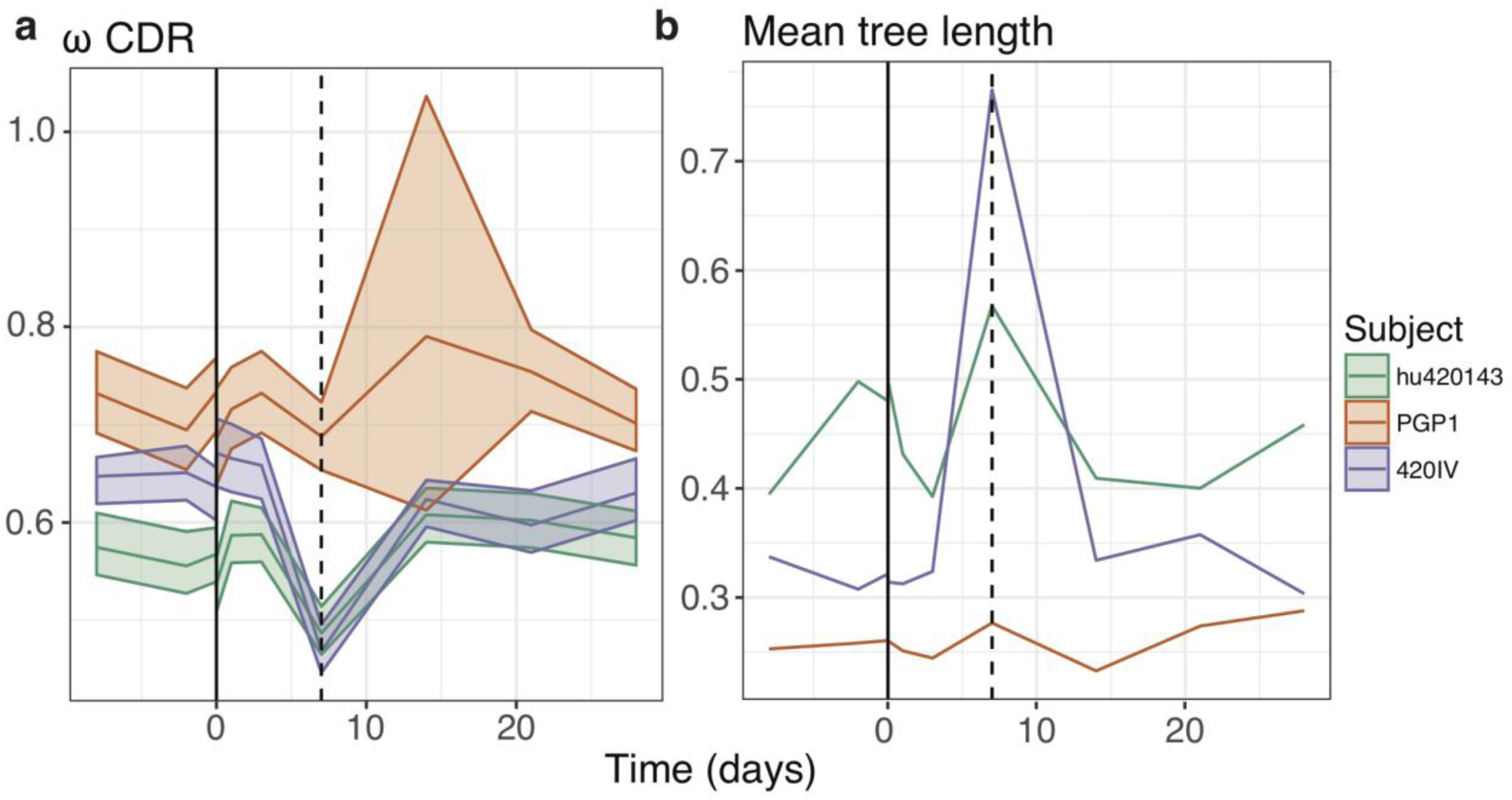
Signatures of selection and diversity following influenza vaccination (**a**) Estimates of the ω_CDR_ parameter of the HLP19 model over the course of influenza vaccination. (**b**) Estimates of mean tree length (total substitutions per codon within a lineage, averaged across all lineages within a repertoire) over the course of influenza vaccination. The x-axis shows the number of days since vaccination; the vertical dashed line represents day 7 post-vaccination. The shaded areas in panel (a) represent the 95% confidence intervals for each parameter estimate, calculated using profile likelihood curves.

We performed simulation analyses to test whether decreased ω_CDR_ at day+7 in subjects hu420139 and 420IV were due to biases in our parameter estimation procedure (*Methods*, **Supplemental File 6**). None of the 20 simulation repetitions performed using the Vaccine dataset were able to reproduce the observed change in ω_CDR_ at day+7 compared to the pre-vaccination time point (-1h; **Supplemental File 6b** and **6c**), demonstrating that these trends are due to factors besides biases in parameter estimation, given the underlying structure of our datasets and their predicted germline sequences.

### Increased tree length is associated with signatures of negative selection

Our analysis of the Vaccine dataset indicated that, in two subjects, there was a concurrent increase in mean tree length and decrease in ω_CDR_, at day +7 following influenza vaccination. We hypothesized that this relationship between ω_CDR_ and tree length might be more general, and tested this hypothesis using log-linear regression across all 27 subjects of the Age dataset and all 30 samples (10 time points from three subjects) of the Vaccine dataset. Across both datasets we observed a consistent and significant negative relationship between both ω_CDR_ and ω_FWR_, and mean repertoire tree length (i.e. the average expected substitutions per codon site across all lineages within the repertoire; **Figure 5**). This trend was surprisingly similar between datasets, with slopes of linear regressions having overlapping 95% confidence intervals, and was particularly strong in the CDRs. For the Age dataset, the slope of a linear regression of ω_CDR_ against the ln(mean tree length) was −0.24 (95% CI = −0.35, −0.14; p < 6×10^-5^), while for the Vaccine dataset the corresponding slope was −0.26 (95% CI = −0.29, −0.23; p < 4×10^-16^). Overall, these regressions show a 32.1% and 41.4% decrease in ω_CDR_ over the range of mean tree length observed in the Age and Vaccine datasets, respectively. A similar, if weaker, relationship was found between ω_FWR_ and ln(mean tree length) (**Figure 5**; details in caption). This indicates that repertoires with longer (i.e. more diverged) lineages are associated with signatures of increased negative selection, particularly in the CDRs.

**Figure 5:**
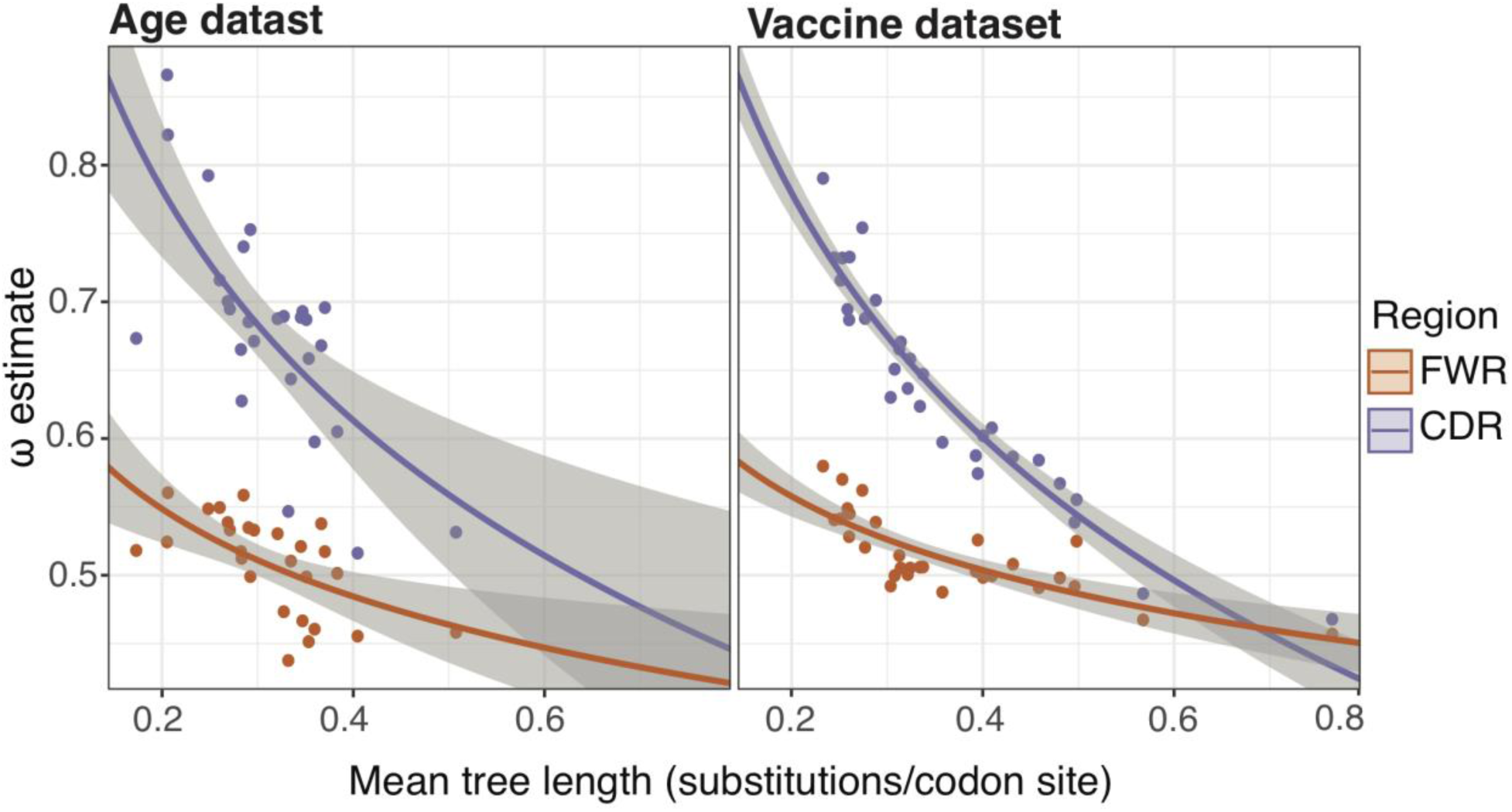
Negative relationship between ω and mean tree length (**a**) Linear regression between estimates of ω_CDR_ (purple) and ω_FWR_ (orange) and the natural log of mean tree length for each subject in the Age dataset. The slope and intercept of ω_CDR_ against ln(mean tree length) were −0.24 (95% CI = −0.35, −0.14) and 0.39, respectively (*p* < 6×10^-5^ for both). The corresponding slope and intercept of ω_FWR_ were −0.09 (95% CI = −0.14, −0.04) and 0.4 (p < 0.002 for both). (**b**) Linear regression between estimates of ω_CDR_ (purple) and ω_FWR_ (orange) and the natural log of mean tree length for each sample in the Vaccine dataset (three subjects, 10 samples each). The slope and intercept of ω_CDR_ against ln(mean tree length) were −0.26 (95% CI = −0.29, −0.23) and 0.36, respectively (*p* < 4×10^-16^ for both). The corresponding slope and intercept of ω_FWR_ were −0.08 (95% CI = −0.1, −0.05) and 0.43 (p < 4×10^-7^ for both). Grey shaded areas in both panels show standard error estimates of the log-linear regression.

We performed simulation analyses to test whether the observed trends between ω and mean tree length were due to biases in our parameter estimation procedure (*Methods*, **Supplemental File 6**). In none of 20 simulations, using both datasets, did we observe a significant relationship between of ω_CDR_ and mean tree length or ω_FWR_ and mean tree length (**Supplemental File 6d-e)**. However, the simulations in **Supplemental File 6** were performed under a fully-context dependent version of the HLP19 model, which does not completely represent the biased nature of SHM. To test whether a richer model of SHM could potentially reproduce our results, we performed simulations using the S5F model (Yaari *et al.* 2013), using the tree topologies and branch lengths estimated using maximum parsimony (dnapars v3.679; Felsenstein 2002) and the predicted germline sequences of the Age dataset (detailed in **Supplemental File 7**). None of 50 such simulation repetitions showed a negative slope between ω_CDR_ and mean tree length as large as that observed for the empirical data; only one simulation repetition showed a more negative slope than observed between ω_FWR_ and mean tree length (**Supplemental File 7**). We therefore conclude that the negative relationship between mean tree length and ω_CDR_ observed in **Figure 5** is not due simply to inherent biases in our parameter estimation procedure.

## Discussion

Phylogenetic techniques have been used to study B cell lineages for many years (Shlomchik *et al.* 1987) and continue to be a powerful tool in understanding affinity maturation (Vieira *et al.* 2018). Two fundamental issues that arise from the application of phylogenetic techniques to B cell repertoires are (i) the biology underlying B cell affinity maturation violates key assumptions of most phylogenetic models, and (ii) phylogenetic models are designed typically to work on lineages originating from a common ancestor, but B cell repertoires are composed of multiple lineages with separate ancestries, many of which are composed of only a few unique sequences. If such lineages are each analyzed independently then parameter estimates will be noisy and highly uncertain. Here we introduce a hierarchical approach to B cell phylogenetics that addresses these issues. We extend phylogenetic models of SHM evolution so that they are consistent with the known biology of B cell affinity maturation, and can share parameters of the sequence evolution process across all lineages within a repertoire. This approach outperforms the alternative of estimating parameters for each lineage individually. By applying our new approach to empirical data we find evidence consistent with dysregulation of SHM in older subjects, increased negative selection during influenza vaccine response, and a relationship between mean tree lengths and signatures of negative selection.

We first used our hierarchical framework to demonstrate a negative association between age and the estimated mutability of W<u>A</u> hot-spot motifs in males and females. Previous studies have shown that aging is associated with a decrease in the affinity, specificity and diversity of antibodies produced (LeMaoult *et al.* 1997; Dunn-Walters *et al.* 2003; Weiskopf *et al.* 2009; Dunn-Walters 2016), as well as with a number of changes at the repertoire level, including longer CDR3s, higher levels of SHM, and persistent clonal lineages in the blood (Wang et al. 2013). It is possible that age-related dysregulation of SHM machinery plays a role in phenomena associated with immunosenescence. Our finding that older individuals tend to have altered mutability of W<u>A</u> motifs is consistent with this hypothesis. However, we note that this effect is observed only in a small cohort, and while the linear regression slopes we observed were significant, the overall change in W<u>A</u> mutability was modest. While we were unable to reproduce this relationship in simulations under a null model (**Supplemental File 6**), it is possible this trend is driven by other confounding factors we have not considered. Future analyses with more subjects are needed to validate this trend generally.

We further used our hierarchical phylogenetic approach to characterize BCR molecular evolution during vaccination. B cell receptors during affinity maturation are subject to multiple selective pressures; positive selection to introduce new affinity-increasing amino acid variants (higher ω) and negative selection to remove affinity-decreasing variants (lower ω). *A priori*, we might expect positive selection to predominate during vaccine response. However, in our analysis we found that lineages present at the time of peak influenza vaccine response show signs of increased negative selection (lower ω) on CDRs. We suggest this is because B-cell lineages with a history of affinity maturation during influenza infection/vaccination will likely have already evolved effective or nearly effective neutralization at the time of vaccination, resulting in a greater proportion of amino acid changes being deleterious (Clarke *et al.* 1985). This would result in lower ω_CDR_ values, which may be particularly marked during influenza vaccine response, since B cells activated by influenza vaccination in adults are expected to derive from re-activated memory B cell lineages (Vollmers *et al.* 2013).

We also observed a negative relationship between tree length and ω_CDR_ across both of our datasets (**Figure 5**). This relationship is remarkably consistent given that our combined datasets contain a total of 30 subjects of different age, sex, and treatment status. None of the simulation analyses performed under a null model were able to reproduce this result (**Supplemental File 6** and **7**), so this relationship is unlikely to be due to a bias or intrinsic correlation between these variables in our estimation procedure. In the absence of other obvious confounding factors, we posit a simple biological explanation: as B cell clones accumulate mutations through repeated rounds of affinity maturation, their binding affinity to target antigen increases and consequently the benefit of random amino acid changes (i.e. new mutations) decreases (Clarke *et al.* 1985). This idea that the rate of fitness-increasing mutations decreases is a straightforward implication of a population nearing a “peak” within a fitness landscape (Wright 1932). Sheng *et al.* (2016) demonstrated evidence of this process (which they termed the “affinity maturation selection” model) in anti-HIV broadly neutralizing antibody (bnAb) lineages. This explanation is also consistent with the findings of Yaari *et al.* (2015) showing that mutations earlier in B cell lineage trees from healthy subjects show clearer signs of positive selection than more recent mutations. Our results suggest that decreased rates of non-synonymous mutations relative to synonymous mutations, as observed in HIV bnAb lineages (Sheng *et al.* 2016), BCR repertoires during vaccine response (**Figures 4** and **5**), and even in healthy subjects with no obvious signs of infection (**Figure 5**; Yaari et al. 2015), are all special cases of a general feature of affinity maturation.

There are several limitations to the hierarchical phylogenetic approach introduced here. The most obvious is that by constraining parameter values so that they are identical for all lineages within a repertoire, we mask any potential parameter variation among lineages. Estimating separate parameter values for one or a set of lineages within a repertoire may be easily accommodated within the hierarchical framework (Suchard *et al.* 2003). This may be useful for parameters such as ω_CDR_, which might reflect lineage-specific histories of antigen-driven selection (Horns *et al.* 2019). As an example of this approach, we explore heterogeneity in estimates of ω_CDR_ among lineages of different sizes for one repertoire (**Supplemental File 9**). It is unclear whether estimation of individual κ and *h*^*a*^ values would yield useful insights, since these parameters relate primarily to biases resulting from SHM, and there is little *a priori* reason to believe they might vary among B cell lineages within an individual. However, it is clear that estimating parameters (e.g. ω_CDR_) for each lineage individually will lead to issues with over-fitting (e.g., when all CDR mutations within a lineage are non-synonymous). Further work will be needed to resolve lineage heterogeneity within individual repertoires.

A hierarchical phylogenetic approach to BCR phylogenetics is justified theoretically and represents a step forward in the statistical analysis of B cell repertoires. Our new methods are implemented in the program IgPhyML (v1.0.7; https://bitbucket.org/kbhoehn/igphyml), which is freely available and integrated into the Immcantation suite (http://immcantation.org).

