## Supplemental Files for "Repertoire-wide phylogenetic models of B cell molecular evolution reveal evolutionary signatures of aging and vaccination"

**Supplemental File 1: Supplemental Methods**

*HLP19 model definition*

We begin with the HLP17 substitution model (Hoehn *et al.* 2017), in which instantaneous rates of codon substitution are parameterized by the nonsynonymous/synonymous mutation rate ratio (ω), transition/transversion mutation rate ratio (κ), codon frequencies (**π**), and modified substitution rates $h^a$ where $a$ is an SHM hot- or cold-spot motif, such as WR<u>C</u> (Peled *et al.* 2008); W=A/T, R=A/G). We modify the HLP17 model by removing codon frequencies from the **Q** matrix, and parameterize the instantaneous rate of change from codon *i* to *j* as:

$$q_{ij} = \begin{cases} 0 & i \to j \; more\ than\ 1\ nucleotide\ change \\ 1 + \sum_a b_{ij}^a h^a & i \to j \; synonymous\ transversion \\ \kappa\left(1 + \sum_a b_{ij}^a h^a\right) & i \to j \; synonymous\ transition \\ \omega\left(1 + \sum_a b_{ij}^a h^a\right) & i \to j \; nonsynonymous\ transversion \\ \omega\kappa\left(1 + \sum_a b_{ij}^a h^a\right) & i \to j \; nonsynonymous\ transition \end{cases}$$

where $b_{ij}^a$ is the probability that a mutation from *i* to *j* involves the underlined base in motif *a* (e.g. WR<u>C</u>, where W = A/T, R = A/G, and C experiences an altered substitution rate). The values of $b_{ij}^a$ are calculated by marginalizing over all possible 5’ and 3’ flanking sense codons as follows:

$$b_{ij}^{a} = \sum_{k=1}^{61}\sum_{g=1}^{61} \pi_k \, \pi_g \begin{cases} 1 & kig \rightarrow kjg \; is \; a \; mutation \; from \; motif \text{ a} \\ 0 & otherwise \end{cases}$$

Importantly, though **π** values are not used in the substitution matrix, **Q** does depend on **π** through **B** and through matrix calibration (see next section).

*Rate matrix calibration*

**Q** matrices are typically calibrated using equilibrium frequencies such that branch lengths are equal to the expected number of substitutions per site (Lio and Goldman 1998). Namely, the mean value of a **Q** matrix (μ) is calculated as:

$$\mu = -\sum_{i=1}^{61} \pi_i q_{ii}$$

Each element in the **Q** matrix is then divided by μ, giving the **Q** matrix a mean value of 1, thus making phylogenetic branch lengths interpretable as the expected number of substitutions per site, assuming state frequencies have converged to a stationary distribution. State frequencies (**π**) are an important part of this calculation, and if they do not accurately reflect the composition of the sequence at a particular time point this may lead to inaccurate branch length estimates. Typically, codon frequencies are assumed to have reached their stationary distribution, and **π** is set to the equilibrium codon frequency distribution. This assumption is made in virtually all substitution models, including HLP17 (Hoehn *et al.* 2017).

Calibrating **Q** matrices using equilibrium frequencies is likely a poor choice for B cell lineages because B cells are known to begin out of sequence equilibrium and change over time (Sheng *et al.* 2016). Because the germline sequence of B cell lineages is predicted before phylogenetic estimation, it is possible to use either the empirical codon frequencies of the germline sequence ($\boldsymbol{\pi^g}$) or the empirical codon frequencies of the tip sequences ($\boldsymbol{\pi^t}$) to calibrate the **Q** matrix. However, both of these reflect codon frequencies at only the beginning and end of the evolutionary process. We instead calibrate our **Q** matrix using codon frequencies at the midpoint of the tree ($\boldsymbol{\pi^m}$). To predict these values, we first calibrate the **Q** matrix using $\boldsymbol{\pi^g}$ to give $\mathbf{Q^g}$. and calculate the transition probability matrix $P^m = \exp(Q^g m)$, where $m$ is half the mean root-tip divergence of all sequences in the initial tree – which we estimate using the GY94 (Goldman and Yang 1994; Nielsen and Yang 1998) model on the same data. From these pieces of information, we calculate $\boldsymbol{\pi^m}$ using:

$$\pi_i^m = \sum_j \pi_j^g P_{ji}^m$$

and use these values to calibrate **Q** to give $\mathbf{Q^m}$, which is used for all likelihood calculations and is scaled such that branch lengths are equal to the expected number of codon substitutions per site at the middle of the starting tree. In addition to branch lengths, $b_{ij}^a$ values also depend on codon frequencies. To solve for these, we use $\boldsymbol{\pi^g}$ values to create the **B** matrix used in constructing $\mathbf{Q^g}$. Ideally, we would then use the estimated $\boldsymbol{\pi^m}$ values to construct the B matrix used in $\mathbf{Q^m}$. However, doing so would require recalculating all $b_{ij}^a$ values each time any parameters is updated. This is computationally expensive, requiring $61^4$ computations rather than $61^2$ computations for simply updating the **Q** matrix. We instead calculate each element of $b_{ij}^a$ once using the values of $\boldsymbol{\pi^g}$ and keep the **B** matrices constant throughout remaining parameter estimation. This dramatically improves runtime. In this way, $\mathbf{Q^m}$ depends on $\boldsymbol{\pi^m}$ through calibration and on $\boldsymbol{\pi^g}$ through $\mathbf{B}$ matrix values.

Further, in many of our analyses, we partition our sequences into complementarity determining regions (CDRs) and framework regions (FWRs), and estimate separate values of $\omega$ for each. In these cases, $\boldsymbol{\pi^g}$ and $\boldsymbol{\pi^m}$ estimation is the same as described above but performed on the separate partitions independently: each tree has two $\mathbf{Q}$ matrices, which are identical except that $\mathbf{Q}_{CDR}^{m}$ uses $\omega_{CDR}$ and is calibrated using codon frequencies in the CDRs, while $\mathbf{Q}_{FWR}^{m}$ uses $\omega_{FWR}$ and is calibrated using codon frequencies in the FWRs. Likelihood calculations involving ambiguous codons in the root sequence are averaged over all compatible codons, weighted by their frequencies in either CDRs or FWRs obtained from an empirical dataset of four healthy subjects (Heiden *et al.* 2017).

Overall, compared to HLP17, this updated model, which we refer to as HLP19, uses less than half the number of free parameters and more reliably models the non-equilibrium nature of affinity maturation. It is also more formally similar to previous nonreversible phylogenetic model forms (Yang 1994; Kaehler *et al.* 2017).

*Repertoire dataset processing*

The "Age" dataset consists of samples taken from 27 healthy individuals in two consecutive years (Wang *et al.* 2013). Subjects varied in age from 20 to 81 years old, and both male and female subjects were included. Sequencing was performed in three replicates: two using genomic

DNA and one using mRNA. Raw bulk-sequencing reads using the 454 (Roche) platform from PBMCs as described in (Wang *et al.* 2013) were preprocessed using pRESTO v0.4.1 (Vander Heiden *et al.* 2014). Briefly, reads were quality controlled by filtering reads with length shorter than 250 nucleotides and those with a mean Phred quality score below 20. Reads were aligned against V and C region primers, allowing for a maximum error rate of 0.2. Duplicate reads and reads that were observed only once (singletons) were removed. Following preprocessing, V(D)J assignment was performed using IgBLAST v1.7.0 (Ye *et al.* 2013) against the IMGT human germline reference database (IMGT/GENE-DB v3.1.16; retrieved June 19th, 2017) (Giudicelli *et al.* 2005). Default IgBLAST parameters were used. Postprocessing of IgBLAST output and clonal clustering were performed using Change-O v0.3.7 (Gupta *et al.* 2015b). Putative non-productively rearranged sequences were filtered. To infer clonal lineages, sequences were first partitioned based on common IGHV gene annotation, IGHJ gene annotation, and junction region length. IGHV and IGHJ gene annotations for each partition were determined by the union of ambiguous assignments within each junction length partition having at least one overlapping gene annotation among all gene assignments. Within these larger groups, sequences differing from one another by a normalized Hamming distance of 0.1 within the junction region were clustered as clones via single-linkage hierarchical clustering (Gupta *et al.* 2017). The clonal distance threshold was determined by manual inspection to identify the local minima between the two modes of the bimodal distance-to-nearest histogram produced using SHazaM v0.1.8)(Gupta *et al.* 2015b). The V and J genes of unmutated germline ancestors of each clone were predicted using Change-O, with D segment and N/P regions masked by "N" nucleotides. Clonal clustering was performed using all three sequencing replicates (two obtained from genomic DNA and one mRNA) pooled together; however, after this mRNA sequences were separated and exclusively used for subsequent analysis.

The “Vaccine” dataset consists of samples from three male donors at 10 time points: -8 days, -2 days, -1 hour, +1 hour, +1 day, +3 days, +7 days, +14 days, and +28 days relative to seasonal influenza vaccination (Laserson *et al.* 2014). The sample collection protocol is available in (Laserson *et al.* 2014); sequencing, as well as data preprocessing for read quality and inference of clonal clusters is described in (Gupta *et al.* 2017). Briefly, samples were re-sequenced from mRNA using 5’RACE with unique molecular identifiers (UMIs) and sequenced using the Illumina MiSeq platform. Using pRESTO (v 0.4; Vander Heiden *et al.* 2014), low quality bases in reads (Phred score < 5) were masked with Ns, paired-end reads were matched and assembled, and assembled sequences with the same UMI were collapsed into a consensus sequence (Gupta *et al.* 2017). Following preprocessing, V(D)J assignment was performed using IgBLAST v1.4.0 (Ye *et al.* 2013) against the IMGT human germline reference database (IMGT/GENE-DB v3.1.16; retrieved June 19th, 2017) (Giudicelli *et al.* 2005). Default IgBLAST parameters were used. Putative non-productively rearranged sequences were filtered. As with the Age dataset, clonal lineages were inferred by grouping sequences based on common IGHV gene, IGHJ gene, and junction length. Within these groups, sequences differing from one another by a normalized Hamming distance of 0.1 within the junction region were clustered into clones via single-linkage hierarchical clustering (Gupta *et al.* 2017). The V and J genes of the unmutated germline ancestors for each clone were predicted using Change-O (v0.4.3; Gupta *et al.* 2015), with D segment and N/P regions masked by ‘N’ nucleotides.

For HLP17/19 phylogenetic analysis it is necessary for all sequences within a particular clone to be placed in a codon alignment with the correct reading frame. To achieve this, sequences from each clone were multiple aligned using the IMGT numbering scheme, which was also used to determine the locations of CDRs and FWRs (Lefranc and Lefranc 2001). It is possible that BCRs will accumulate insertion mutations relative to their germline precursor over the course of affinity maturation. In this case, Change-O removes insertions in its IMGT sequence alignment in order to preserve numbering. However, frame-preserving insertions occasionally occur mid-codon. Removing these insertions brings together nucleotides that do not actually form an observed codon in the sequence, creating a potentially false codon substitution. We detected these events by comparing input sequences to IMGT aligned sequences in Change-O files, and masking them using "N" nucleotides. We then trimmed all gap-only columns in IMGT alignments within each clone. This functionality, as well as converting Change-O files to IgPhyML inputs, is now included in the BuildTrees tool as part of Change-O v0.4.3+ (Gupta *et al.* 2015).

Sequences from both years within the same patient were pooled together in the Age dataset, resulting in a single repertoire sample per subject. From both datasets, lineages with fewer than 2 unique sequences (not including the predicted germline) were discarded. Due to the large number of sequences per sample in our Vaccine dataset, these repertoires were subsampled at each subject and time point combination. Sequences from each sample were subsampled without replacement until either the sample contained at least 3,000 unique sequences within clones of at least 2 unique sequences (occasionally this resulted in some datasets having up to 3,001 sequences), or all sequences were used. This cutoff was chosen because all samples from subject hu420139 and 420IV, and 5/10 samples from PGP1 had >3000 unique sequences in non-singleton clones.

**Supplemental File 2: Simulating under the HLP19 model**

We performed multiple simulation analyses to validate our parameter estimation procedure under a fully context-dependent version of the HLP19 model (**Supplemental File 3**, **4**, and **5**). These were performed under different parameter values, tree topologies, and initial sequences as detailed in their respective files, but they share a common means of simulating sequence changes along a given tree. These processes require a starting tree topology, branch lengths, a germline sequence, and substitution parameters to be specified in advance. For a given lineage tree, the starting germline progenitor sequence is specified as either a random naïve B cell sequence (**Supplemental File 3**) or the previously identified germline sequence in the empirical data (see **Supplemental Files 4** and **6**). For each branch in the tree we simulated the number of nucleotide substitutions by drawing from a Poisson distribution whose mean equals the branch length divided by three and multiplied by the number of codons in the germline sequence. Branch lengths were divided by three because branches lengths estimated under all substitution models used in this paper are in units of expected codon substitutions per site, while the simulation procedure operates on single nucleotides. For a given sequence we then calculated the probability of all single substitutions under the HLP19 **Q** matrix (**Supplemental File 1**) based on their full nucleotide context (rather than the mean field approximation; described by parameters $h^{WRC}$, $h^{GYW}$, $h^{WA}$, $h^{TW}$, $h^{SYC}$, and $h^{GRS}$), whether the substitution was synonymous or non-synonymous (described by parameters $\omega_{FWR}$ and $\omega_{CDR}$ depending on whether the site was in the FWRs or CDRs), and whether the substitution was a transition or transversion (described by $\kappa$). This process began at the specified germline progenitor and continued down the tree to the tips. We then masked the CDR3 of the germline sequence to mimic the behavior of the

CreateGermlines prediction program, which is unable to reliably predict germline D segment and N/P regions (Gupta *et al.* 2015a). We also removed the CDR3 regions of all sequences in each dataset to avoid potential issues generated with our inability to properly predict this region. This process repeated for all clonal lineage trees in a dataset. These simulations were performed using custom Perl scripts, which are available at https://bitbucket.org/kbhoehn/hierarchical-models-scripts.

*Issues with the simulation approach in Hoehn et al 2017*

Hoehn *et al.* (2017) used a similar approach to simulate sequences under the HLP17 substitution model. However, the simulations in Hoehn *et al.* (2017) differ from the approach described above; specifically, in Hoehn *et al.* (2017) the relative substitution probabilities for each site were computed by calibrating the **Q** matrix of the HLP17 model with its codon frequencies so that branch lengths could be interpreted as substitutions per site (discussed in **Supplemental File 1**), and then exponentiating this matrix to give the **P** matrix of substitution probabilities for each codon. Codon substitutions were then introduced based on these probabilities at each site. However, because **Q** matrixes are calibrated using codon frequencies, the degree to which branch lengths can be accurately interpreted as substitutions per site depends on how accurately the codon frequencies used reflect the composition of the sequences being simulated. For the HLP17 model, these frequencies are estimated through maximum likelihood as free parameters – they are not necessarily tied to the composition of the sequences. This results in branch lengths that less accurately reflect the expected substitutions per site than standard models (this inaccuracy is confirmed in **Supplemental File 3**). Unfortunately, because the simulations performed in that paper used calibrated, exponentiated **Q** matrices, this inaccuracy was present in the simulated data itself, leading to simulated datasets with fewer substitutions than expected given their branch lengths. This resulted in the HLP17 model outperforming the GY94 model in branch length estimation when compared to the original branch lengths of the trees used in simulations. While an accurate description of the simulation procedure used, this result is likely specific to this particular simulation approach and not general, and does not hold up in the fairer trials presented in this paper. The approach taken here (outlined above) avoids this issue by simply weighting all possible single nucleotide substitutions for all sites in a sequence by their values in the substitution matrix, randomly choosing a substitution, and then repeating this process on the new sequence for a given number of substitutions. This simulation process makes no explicit assumptions regarding codon frequencies – they change as a result of the substitution parameters, starting sequence, and lineage tree length. Further, the simulation analyses used in this paper explicitly define the number of nucleotide substitutions added on each branch. This means branch length estimates in the analyses presented in this paper (**Supplemental File 3**) are scored for accuracy based on the actual number of substitutions per codon site used in the simulation procedure, rather than the branch lengths of the original tree topology as done in Hoehn *et al.* (2017).

**Supplemental File 3: Performance of GY94, HLP17, and HLP19 substitution models**

The goal of this simulation analysis is to compare the parameter estimation performance of the GY94 (Goldman and Yang 1994; Nielsen and Yang 1998), HLP17 (Hoehn *et al.* 2017), and HLP19 substitution models. Simulations were performed on two trees with identical topologies and total tree length (i.e. expected number of substitutions), but different branch length distributions (*split* and *long*; see **Supplemental Figure 3a** below). We created artificial datasets by simulating sequences using 200 copies of either *split* or *long* tree topologies. Simulations were performed as described in **Supplemental File 2**. In these analyses, we selected a random naïve heavy chain sequence from a dataset of two healthy individuals (Heiden *et al.* 2017) for each tree in each dataset. We further used two sets of substitution parameters. In the first, named *hot*, $\kappa = 2$, $\omega_{FWR} = 0.4$, $\omega_{CDR} = 0.7$, $h^{WRC} = 2$, $h^{GYW} = 3$, $h^{WA} = 2$, $h^{TW} = 1$, $h^{SYC} = -0.6$, and $h^{GRS} = -0.6$. In the second, *hotter*, $\kappa = 2$, $\omega_{FWR} = 0.4$, $\omega_{CDR} = 0.7$, $h^{WRC} = 4$, $h^{GYW} = 6$, $h^{WA} = 4$, $h^{TW} = 2$, $h^{SYC} = -0.6$, and $h^{GRS} = -0.6$. Overall, we created 50 datasets for each combination of tree topology and substitution model.

To compare model performance, for each simulated dataset we used maximum likelihood to estimate tree topology, branch lengths, $\kappa$, $\omega_{FWR}$, and $\omega_{CDR}$ the GY94, HLP17 and HLP19 models. We also estimated $h^{WRC}$, $h^{GYW}$, $h^{WA}$, $h^{TW}$, $h^{SYC}$, and $h^{GRS}$ for HLP17 and HLP19. All of these parameters were estimated at the hierarchical level. Codon frequencies ($\pi$) shared among all lineages within the repertoire were estimated using maximum likelihood for the HLP17 model, as empirical frequencies within each lineage (pooled across FWR and CDRs) for GY94, and using the procedure detailed in **Supplemental File 1** for HLP19. The CDR3 region was removed for all sequences after simulation. For each parameter, we calculated bias as the mean (true–estimated)/true value across all 50 datasets of each type. True branch lengths for each simulated lineage were defined as the number of substitutions that occurred outside of the CDR3 along a particular branch, divided by the number of codon sites outside of the CDR3 in the simulated alignment. The results of these tests are shown in **Supplemental Figure 3b** and **c** below.

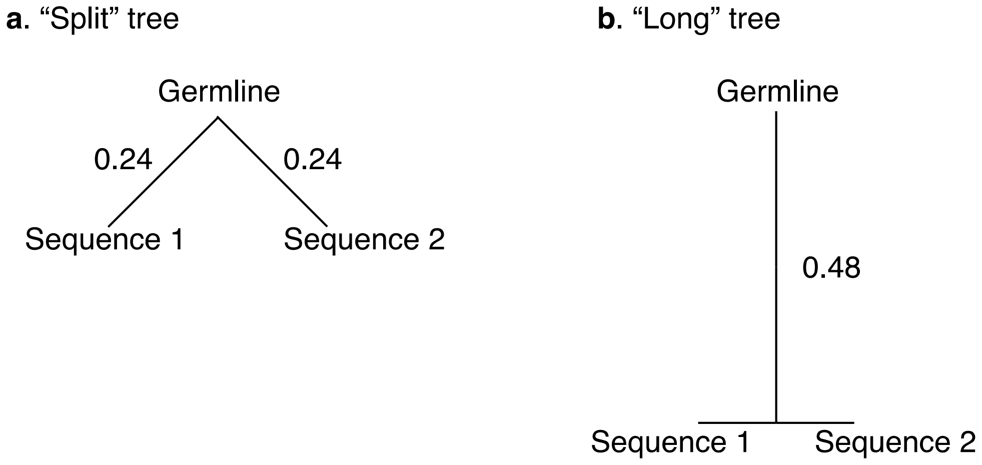


*Supplemental Figure 3a*: Trees used in simulation analyses described in **Supplemental File 3**. Repertoires specified as "split" used the tree in **a**, and repertoires specified as "long" used the tree in **b**. Branch lengths are in codon substitutions per site. All branches with unlabeled lengths are equal to zero. "Germline" is the germline progenitor sequence of the lineage, while Sequence 1 and Sequence 2 are the data sequences produced by the simulation process.

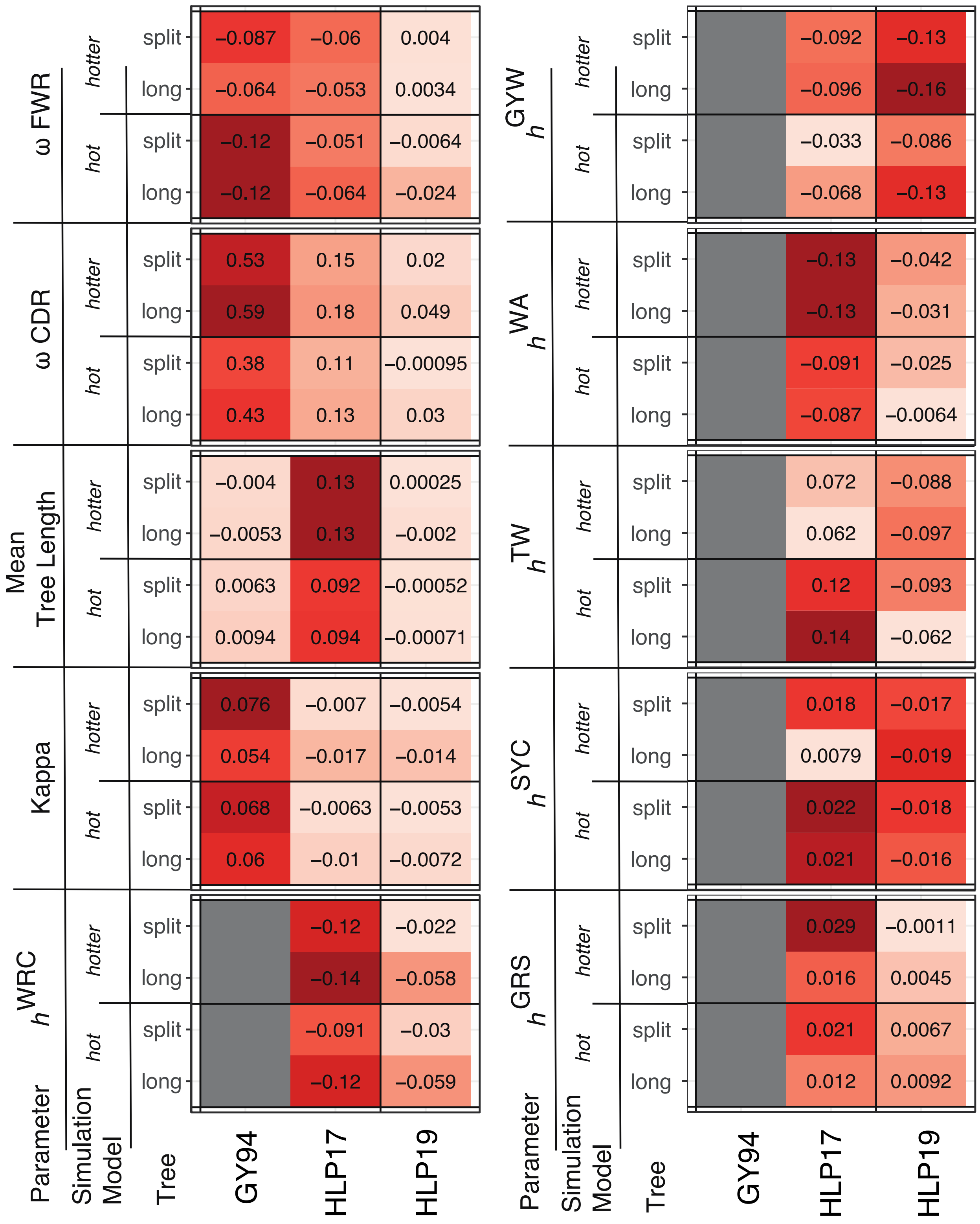


*Supplemental Figure 3b*: Mean bias in parameter estimation for different substitution models. Y axis labels show the parameter estimated, as well as the tree type and model used in simulation. X axis labels show the substitution model used to estimate parameters. See **Supplemental File 3** for full details.

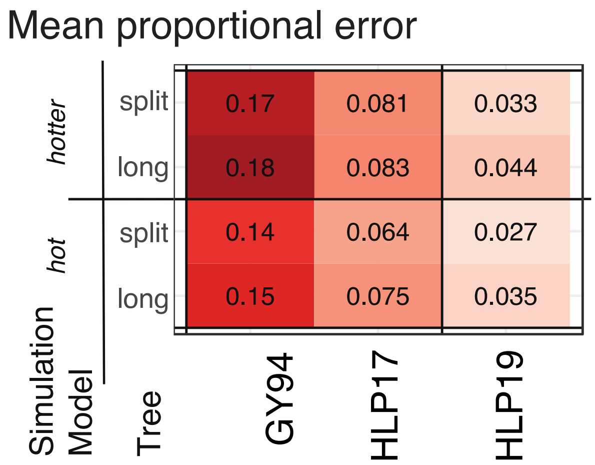


*Supplemental Figure 3c*: Mean absolute bias in substitution parameter estimation for different substitution models. Mean error was calculated as the mean absolute value all parameters shown in Supplemental Figure 3b. *h* values were not measured for GY94, so were not included in this calculation.

**Supplemental File 4: Performance of individual vs hierarchical models in parameter estimation**

The goal of this simulation analysis was to quantify improvement in precision of the hierarchical phylogenetic approach vs individual lineages using the HLP19 substitution model. We used lineage trees from a single repertoire - subject 97 in the Age dataset - which was chosen due to its greater number of large lineages compared to other samples in the Age dataset. As described in **Methods**: *Phylogenetic model parameter and topology estimation*, we first estimated maximum likelihood tree topologies, branch lengths, and shared values of $\omega$, and $\kappa$ for all lineages within each repertoire under the GY94 model. Codon base frequencies were set to their empirical estimates across all lineages within the repertoire using a CF3X4 model (Kosakovsky Pond *et al.* 2010). For each lineage tree within each repertoire, we used the predicted germline sequence (see *Repertoire dataset processing* in **Supplemental File 1**) of the empirical data as the progenitor of each lineage. Masked codons (specified by 'NNN') were replaced with codons drawn from a random uniform distribution. These were primarily in the CDR3 region and were removed after simulations. Simulations were then performed using the procedure outlined in **Supplemental File 2**, using substitution parameters $\kappa = 2$, $\omega_{FWR} = 0.5$, $\omega_{CDR} = 0.7$, $h^{WRC} = 4$, $h^{GYW} = 6$, $h^{WA} = 3$, $h^{TW} = 1$, $h^{SYC} = -0.6$, and $h^{GRS} = -0.6$. These were chosen to be similar to those estimated from the Age dataset (**Figure 1**). We simulated 5 repertoire datasets from the Age and Vaccine datasets using this procedure and removed the CDR3 regions of all simulated sequences. We then repeated the parameter estimation procedure in **Methods**: *Phylogenetic model parameter and topology estimation* by (briefly) first estimating tree topologies, branch lengths, and values of $\omega$, and $\kappa$ under the GY94 model, and then estimating substitution parameters under the HLP19 model. For GY94 estimations, codon base frequencies were set to their empirical estimates across all lineages within the repertoire using a CF3X4 model (Kosakovsky Pond *et al.* 2010). We repeated this process both for each lineage individually, and for each repertoire using a hierarchical model. For individual trees, all parameters were estimated for each individual lineages. For repertoire-wide parameter estimates, all parameters were uniform among all lineages within a repertoire. These results are shown in **Supplemental Table 4** below.

| Parameter estimation | Individual, all lineages | Individual, ≥10 sequences | Repertoire-wide |
|---|---|---|---|
| $\kappa$ | 5.42 | 0.23 | **0.03** |
| $\omega_{FWR}$ | 26.19 | 0.23 | **0.10** |
| $\omega_{CDR}$ | 211.35 | 25.68 | **0.04** |
| $h^{WR\underline{C}}$ | 5.44 | 0.44 | **0.10** |
| $h^{\underline{G}YW}$ | 3.37 | 0.38 | **0.13** |
| $h^{W\underline{A}}$ | 5.99 | 0.42 | **0.04** |
| $h^{\underline{T}W}$ | 10.00 | 0.87 | **0.08** |
| $h^{SY\underline{C}}$ | 3.99 | 0.70 | **0.03** |
| $h^{\underline{G}RS}$ | 8.19 | 0.81 | **0.12** |
| Mean error | 33.97 | 3.62 | **0.07** |

*Supplemental Table 4*: Mean absolute proportional error of individual and hierarchical estimates of HLP19 model parameters from 289 lineages simulated for five repetitions under the HLP19 substitution model. All lineages had ≥2 unique sequences, 256 had < 10 sequences, 33 had ≥ 10 sequences.

### Supplemental File 5: Comparison of model fit on empirical datasets

The Akaike information criterion (AIC) is means of scoring the quality of model fit to data that compares the number of freely estimated parameters to the maximum log likelihood calculated using the model (Akaike 1974). Lower AIC values indicate a better model fit. Here, we calculate AIC for GY94 (Goldman and Yang 1994; Nielsen and Yang 1998), HLP17 (Hoehn *et al.* 2017), and HLP19 (**Supplemental File 1**) models when our parameter estimation procedure (**Methods**: *Phylogenetic model parameter and topology estimation*) was applied to each subject of the Age dataset (**Supplemental Table 5**). Because GY94 and HLP17 are expected to overestimate $\omega_{CDR}$ (**Supplemental File 3**), we also report ML estimates of $\omega_{CDR}$ for each model. In all subjects, both AIC and $\omega_{CDR}$ followed the pattern GY94 > HLP17 > HLP19, indicating HLP19 had both a better fit to the datasets and lower estimates of $\omega_{CDR}$.

The GY94 model used 3 free parameters ($\kappa$, $\omega_{FWR}$, and $\omega_{CDR}$), HLP17 used 18 ($\kappa$, $\omega_{FWR}$, $\omega_{CDR}$, $h^{WRC}$, $h^{GYW}$, $h^{WA}$, $h^{TW}$, $h^{SYC}$, $h^{GRS}$, and three free base frequencies for each of three codon sites under the CF3X4 model (Kosakovsky Pond *et al.* 2010), and HLP19 used 9 ($\kappa$, $\omega_{FWR}$, $\omega_{CDR}$, $h^{WRC}$, $h^{GYW}$, $h^{WA}$, $h^{TW}$, $h^{SYC}$, $h^{GRS}$). To make AIC values comparable among the three models, we altered the HLP17 and HLP19 models slightly by multiplying the partial likelihood of each possible codon at the root by the frequency of that codon ($\boldsymbol{\pi}$), as is typically done for reversible models (Felsenstein 1981). For GY94 these frequencies were the empirical estimates of codon frequencies under a CF3X4 model for each lineage within the repertoire, for HLP17 these frequencies were estimated by ML using the CF3X4 model and shared across all lineages within the repertoire, while for HLP19 these frequencies were estimated from the root sequence of each lineage ($\pi^g$ for FWR or CDR, **Supplemental File 1**).

| | AIC | | | $\omega_{CDR}$ | | |
|---|---|---|---|---|---|---|
| Subject | GY94 | HLP17+$\pi$ | HLP19+$\pi$ | GY94 | HLP17+$\pi$ | HLP19+$\pi$ |
| 1 | 363603 | 360109 | 309005 | 0.99 | 0.77 | 0.69 |
| 3 | 607461 | 602820 | 513667 | 1.04 | 0.80 | 0.70 |
| 4 | 243193 | 241238 | 206345 | 1.03 | 0.83 | 0.74 |
| 8 | 733364 | 727169 | 619748 | 0.99 | 0.77 | 0.67 |
| 11 | 573867 | 567545 | 490976 | 0.86 | 0.67 | 0.60 |
| 13 | 472848 | 469177 | 398199 | 1.14 | 0.88 | 0.79 |
| 14 | 171315 | 170651 | 142884 | 1.00 | 0.77 | 0.67 |
| 15 | 247645 | 245197 | 208959 | 0.79 | 0.60 | 0.55 |
| 16 | 363741 | 360629 | 307737 | 1.07 | 0.86 | 0.75 |
| 18 | 453782 | 448849 | 387236 | 0.91 | 0.68 | 0.60 |
| 21 | 264276 | 261115 | 226276 | 0.79 | 0.59 | 0.52 |
| 23 | 108222 | 107762 | 90691 | 1.18 | 0.93 | 0.87 |
| 25 | 354698 | 351053 | 302490 | 1.00 | 0.78 | 0.69 |
| 26 | 461696 | 457367 | 392300 | 1.04 | 0.78 | 0.69 |
| 33 | 494143 | 488992 | 419978 | 1.02 | 0.78 | 0.69 |
| 34 | 434464 | 430241 | 369880 | 0.93 | 0.72 | 0.64 |
| 49 | 402052 | 398526 | 340069 | 0.98 | 0.77 | 0.69 |
| 54 | 532664 | 526649 | 455190 | 1.01 | 0.79 | 0.67 |
| 63 | 202962 | 201174 | 171439 | 0.95 | 0.72 | 0.63 |
| 64 | 281633 | 279516 | 237513 | 0.99 | 0.78 | 0.69 |
| 65 | 337095 | 333429 | 288345 | 0.99 | 0.76 | 0.69 |
| 73 | 217224 | 215238 | 186094 | 1.06 | 0.85 | 0.70 |
| 74 | 274756 | 272262 | 233127 | 1.06 | 0.79 | 0.67 |
| 83 | 312956 | 309557 | 266105 | 0.99 | 0.75 | 0.66 |
| 93 | 249065 | 247057 | 210139 | 1.09 | 0.83 | 0.72 |
| 95 | 216803 | 215498 | 181918 | 1.22 | 0.95 | 0.82 |
| 97 | 313170 | 308940 | 271512 | 0.77 | 0.61 | 0.53 |

**Supplemental Table 5**: AIC values (rounded to the nearest integer) and $\omega_{CDR}$ estimates of all subjects in the Age dataset using the GY94, HLP17 and HLP19 substitution models, modified as detailed in **Supplemental File 5**.

**Supplemental File 6: Simulation analyses based on empirical datasets**

The goal of these simulation analyses is to test whether differences in the underlying tree topology and branch lengths of samples within the Age and Vaccine dataset could be responsible for observed trends between substitution model parameters and age in the Age dataset and time in the Vaccine dataset. Sequences were simulated in the same manner as in **Supplemental File 4**. As described in **Methods**: *Phylogenetic model parameter and topology estimation*, we first estimated maximum likelihood tree topologies, branch lengths, and shared values of $\omega$, and $\kappa$ for all lineages within each repertoire under the GY94 model. Codon base frequencies were set to their empirical estimates across all lineages within the repertoire using a CF3X4 model (Kosakovsky Pond *et al.* 2010). For each lineage tree within each repertoire, we used the predicted germline sequence (see *Repertoire dataset processing* in **Supplemental File 1**) of the empirical data as the progenitor of each lineage. Masked codons (specified by 'NNN') were replaced with codons drawn from a random uniform distribution. These were primarily in the CDR3 region and were removed after simulations. Simulations were then performed using the procedure outlined in **Supplemental File 2**, using substitution parameters $\kappa = 2$, $\omega_{FWR} = 0.5$, $\omega_{CDR} = 0.7$, $h^{WRC} = 4$, $h^{GYW} = 6$, $h^{WA} = 3$, $h^{TW} = 1$, $h^{SYC} = -0.6$, and $h^{GRS} = -0.6$. These were chosen to be similar to those estimated from the Age dataset (**Figure 1**). We simulated 20 repertoire datasets for each sample in both datasets using this procedure and removed the CDR3 regions of all simulated sequences. We then repeated the parameter estimation procedure in **Methods**: *Phylogenetic model parameter and topology estimation* by first estimating tree topologies, branch lengths, and shared values of $\omega$, and $\kappa$ for all lineages within each repertoire under the GY94 model, and then estimating substitution parameters across all lineages under a hierarchical HLP19 model. For GY94 parameter estimation, codon base frequencies were set to their empirical estimates across all lineages within the repertoire using a CF3X4 model (Kosakovsky Pond *et al.* 2010).

We first investigated whether our simulations could reproduce the observed negative relationship between age, sex, and estimates of $h^{WA}$. For each repetition of our Age dataset of 27 subjects, we fitted a multiple linear regression model with age and sex as interaction variables against substitution rate biases of W<u>A</u> motifs (i.e. $h^{WA}$). We compared the slope coefficients estimated from each of our simulated datasets to those observed in our empirical data. For all 20 repetitions the $h^{WA}$ slope coefficents for males and females were closer to zero than their respective empirical estimates (**Supplemental Figure 6a**).

We next investigated whether our simulations could reproduce the behavior of $\omega_{CDR}$ over the time course of influenza vaccination in our Vaccine dataset. As shown in **Supplemental Figure 6c**, estimates of $\omega_{CDR}$ varied fairly constantly around a mean slightly above the true value (0.7) for the entire time course for all three subjects. For subjects 420IV and hu420139, we calculated the fold change in $\omega_{CDR}$ at day +7 compared to the pre-vaccine sample (-1 hour) for each simulation, and compared it to the empirical estimate. No simulation reproduced the observed fold change at day +7 in either subject (**Supplemental Figures 6b**). As expected given its low sequence count, PGP1 day +14 had wide variation in $\omega_{CDR}$ estimate between repetitions. **Supplemental Figure 6c** confirms that the simulation procedure reliably reproduced differences in mean tree length among repertoire datasets with little variation among repetitions.

We then tested whether our results in either the Age or Vaccine dataset could reproduce our observed negative relationship between ω and mean tree length. For each repetition of our Age and Vaccine datasets, we fit a log-linear regression model in which either $\omega_{CDR}$ or $\omega_{FWR}$ are modelled against the natural logarithm of mean tree length. We compared the slope coefficients estimated from each of our simulated datasets to those observed in our empirical data. For all 20 repetitions, slopes of both ω parameters from both the Age and Vaccine dataset were far lower than any slope parameters observed in the empirical Age or Vaccine datasets (**Supplemental Figure 6d**). These results are further shown in **Supplemental Figure 6e**, which shows these simulation results plotted as regressions and scatterplots.

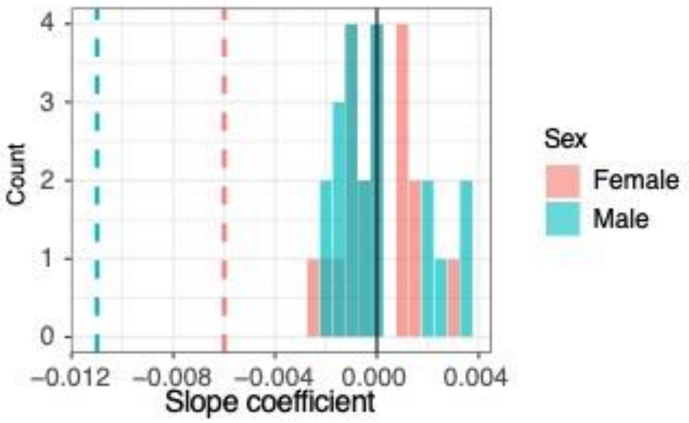


Supplemental Figure 6a: Slope coefficients between $h^{WA}$ and age for 20 null simulation repetitions of the Age dataset. Coefficients are colored by sex. Vertical dashed lines show the values observed in empirical data for their respective parameter and sex.

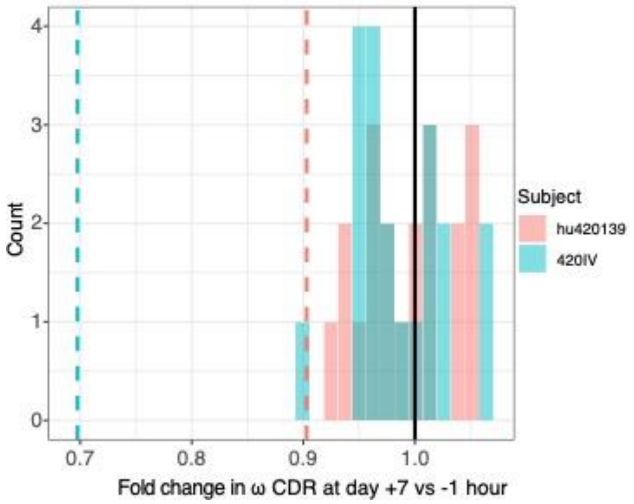


Supplemental Figure 6b: Fold change in $\omega_{CDR}$ between day +7 and -1 hour (pre-vaccination) estimated for 20 null simulation repetitions for subjects hu420139 and 420IV (histograms) and empirical data (vertical lines).

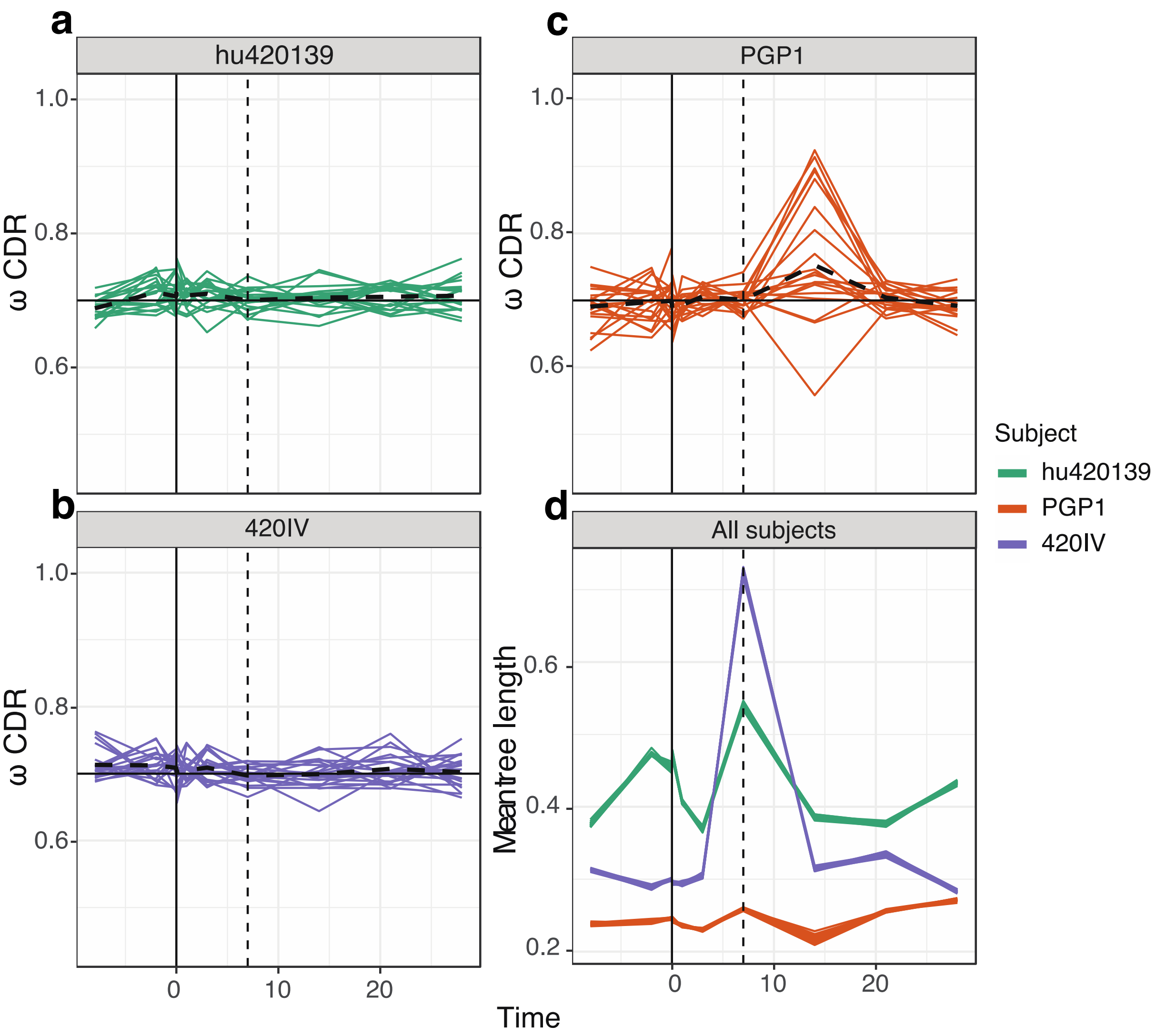


Supplemental Figure 6c: **a-c**: plots of $\omega_{CDR}$ over time for each subject in the Vaccine dataset. Each line represents a repetition of the simulation procedure for a subject. The dashed black line shows the mean value across repetitions at a time point. The true value of $\omega_{CDR}$ in these simulations was 0.7, shown by the solid horizontal black line in each plot. **d**: plots of mean tree length over time for all subjects in the Vaccine dataset. Each line represents a repetition of the simulation procedure for a subject. These lines are closely overlaid due to the lack of variance in this parameter among repetitions.

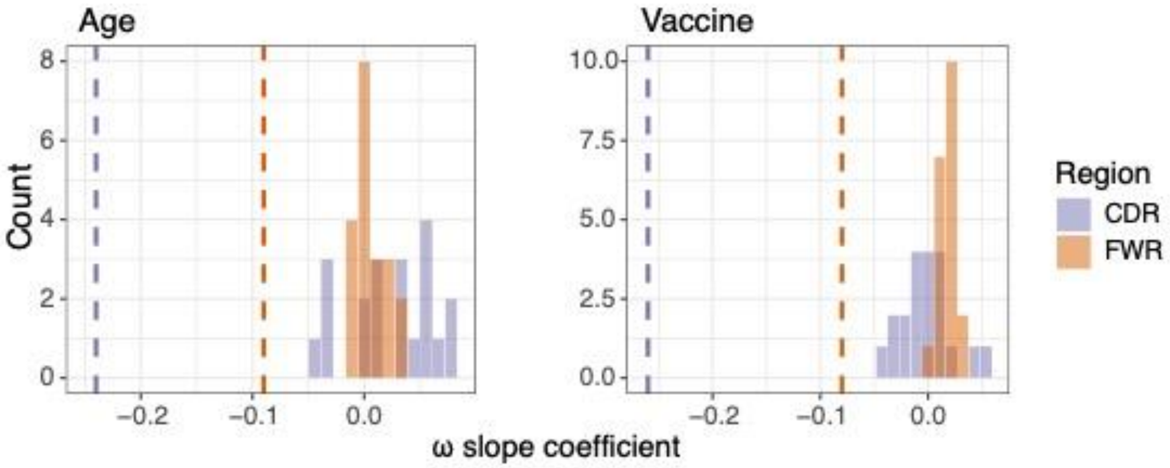


Supplemental Figure 6d: Slope coefficients of log-linear regressions between ω and mean tree length in both the Age and Vaccine datasets among 20 simulation repetitions. Coefficients are colored by region (i.e. whether they represent $\omega_{CDR}$ or $\omega_{FWR}$). Vertical dashed lines show the values observed in their respective empirical datasets.

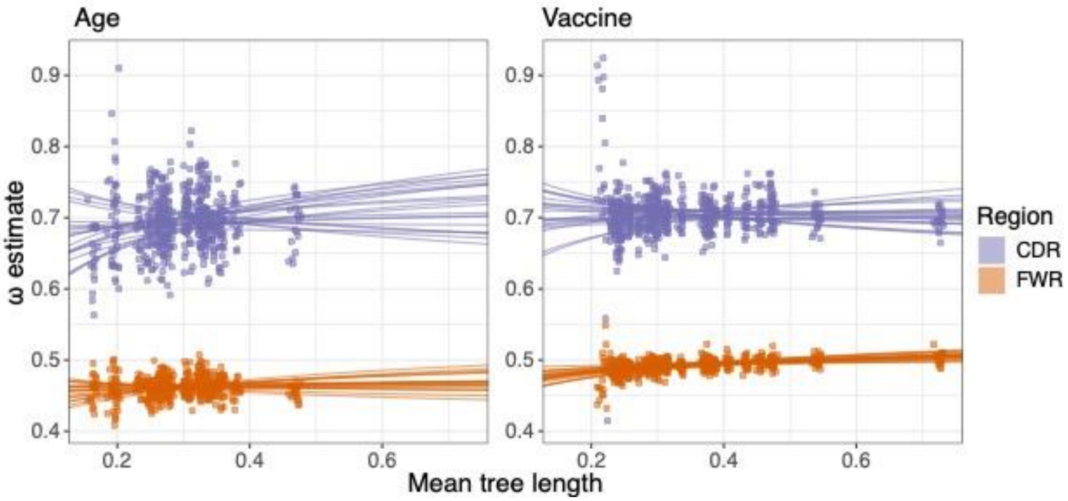


Supplemental Figure 6e: Log-linear regressions between ω and mean tree length among 20 simulation repetitions of the Age and Vaccine datasets. These represent the same data as in **Supplemental Figure 6c**. Coefficients are colored by region (i.e. whether they represent $\omega_{CDR}$ or $\omega_{FWR}$). The true value for all simulations $\omega_{CDR} = 0.7$ and $\omega_{FWR} = 0.5$. The y axis is scaled identically between the two plots.

**Supplemental File 7: Additional simulations using SHMulate**

As an additional validation, we also performed simulations based on the Age dataset using the S5F model within SHazaM (Yaari *et al.* 2013). Simulations were performed under custom versions of Alakazam (v0.2.11; Gupta *et al.* 2015) and SHazaM (v1.9.0; Yaari *et al.* 2013) which are noted below and are available at: https://bitbucket.org/kbhoehn/hierarchical-models-scripts. These simulations for each clone were performed as follows:

1. For a given set of clonal sequences of at least two unique sequences, maximum parsimony lineage tree topology and branch lengths were estimated using *dnapars* from PHYLIP (v3.697; Felsenstein 2002), implemented in Alakazam (v0.2.11; Gupta *et al.* 2015). In contrast to the default behavior in Alakazam of estimating branch lengths using the Hamming distance between reconstructed intermediate sequences, branch lengths were estimated from PHYLIP as the estimated substitutions per site. Trees are also kept strictly binary and do not collapse zero-length branches in internal nodes.
2. Alakazam uses ‘N’ nucleotides to place all sequences within a clone into alignment with respect to IMGT numbering, which adds uninformative sites to the alignment. This means branch lengths taken directly from *dnapars* output – in units of substitutions per site – will be artificially low (this is not a problem in the default setting since *dnapars* branch lengths are not used directly). To correct for this, we re-scale branch lengths; namely, we multiply each branch length by the number of sites in the full alignment, and divide by the number of informative sites (codon sites with any non-N nucleotides in any data sequence) in the alignment.

3. Branch lengths are converted into the number of substitutions between nodes by multiplying each branch length by the number of informative sites in the germline sequence, and randomly rounding up or down based on difference from the nearest integer.
4. In contrast to the default behavior of SHazaM (v1.9.0; Yaari *et al.* 2013), simulations begin with the clone's predicted germline sequence (GERMLINE_IMGT_D_MASK) from CreateGermlines.py from Change-O (v0.4.3; Gupta *et al.* 2015). Mutations are added to the starting sequence based on their relative probability under the S5F model (Yaari *et al.* 2013). Starting from the germline sequence at the root node of the tree, this process is repeated for each descendent node in the tree, with the number of mutations added between ancestor and child nodes equal to the branch length between them.
5. This process is repeated for all clones within a subject in the Age dataset.
6. These simulated datasets are converted into an IgPhyML-readable format using BuildTrees.py in Change-O (v0.4.5; Gupta *et al.* 2015). We then repeat our parameter estimation procedure (see **Methods**:*Phylogenetic model parameter and topology estimation*) by first estimating maximum likelihood tree topologies, branch lengths, and repertoire-wide substitution parameters using the GY94 model, and then using these tree topologies to estimate maximum likelihood branch lengths and substitution model parameters under the HLP19 model.

For each simulation repetition of all 27 subjects, we fit a log-linear regression model in which either $\omega_{CDR}$ or $\omega_{FWR}$ are modelled against ln(mean tree length). We compared the slope coefficients estimated from each of our simulated datasets to those observed in our empirical data. Of the 50 simulation repetitions performed, none showed a slope as negative as that observed between $\omega_{CDR}$ and mean tree length, and only one showed a more negative slope than that observed between $\omega_{FWR}$ and mean tree length (**Supplemental Figure 7a**). These results are further shown in **Supplemental Figure 7b**, which shows these simulation results plotted as regressions and scatterplots. From this we conclude that even under this different simulation framework, which uses a model of SHM not fully captured by the HLP19 model, that the observed trend is not the result of model bias due between tree length and $\omega$, or selection of germline sequence.

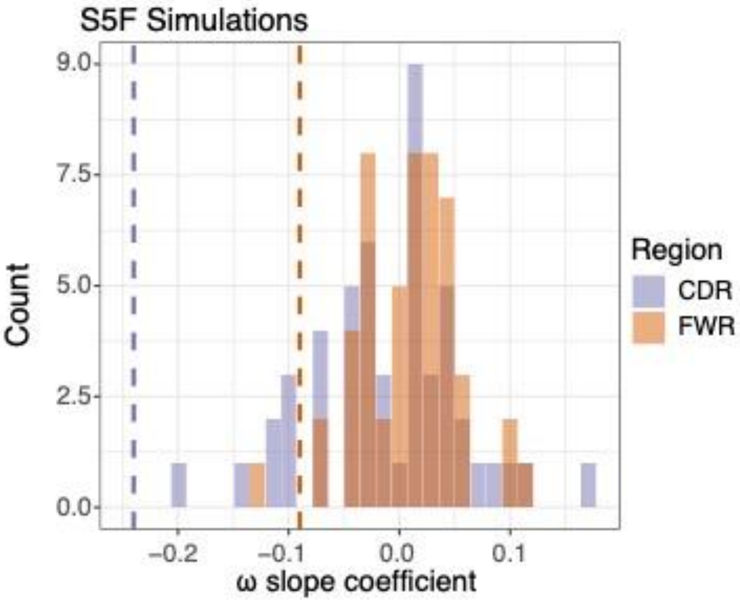


Supplemental Figure 7a: Slope coefficients of log-linear regressions between ω and mean tree length among 50 S5F simulation repetitions of the Age dataset. Coefficients are colored by region (i.e. whether they represent $\omega_{CDR}$ or $\omega_{FWR}$). Vertical dashed lines show the values observed in their respective empirical datasets.

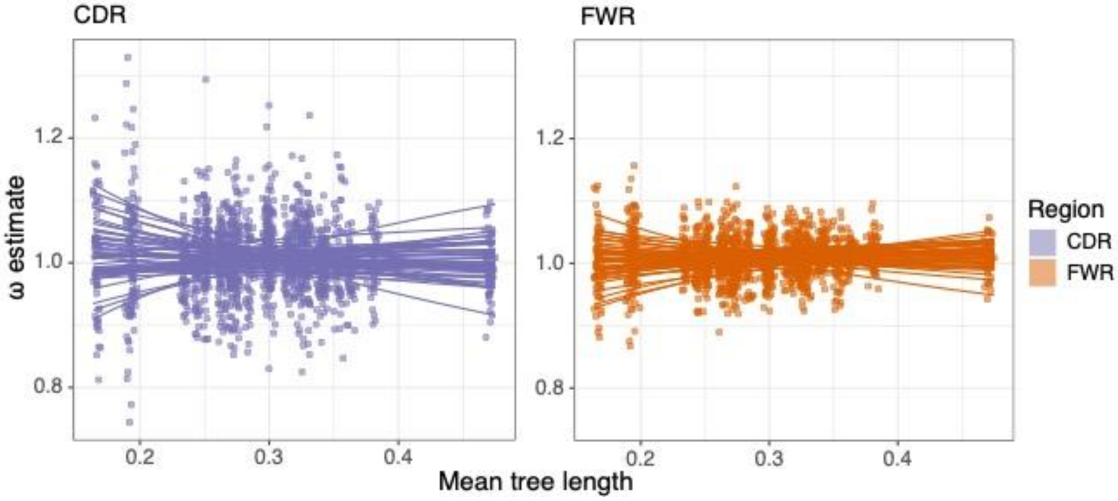


Supplemental Figure 7b: Log-linear regressions between ω and mean tree length among 50 S5F simulation repetitions of the Age dataset. These represent the same data as in **Supplemental Figure 7a**. Coefficients are colored by region (i.e. whether they represent $\omega_{CDR}$ or $\omega_{FWR}$). The Y axis is scaled identically between the two plots.

**Supplemental File 8: Streams plots of clonal sizes**

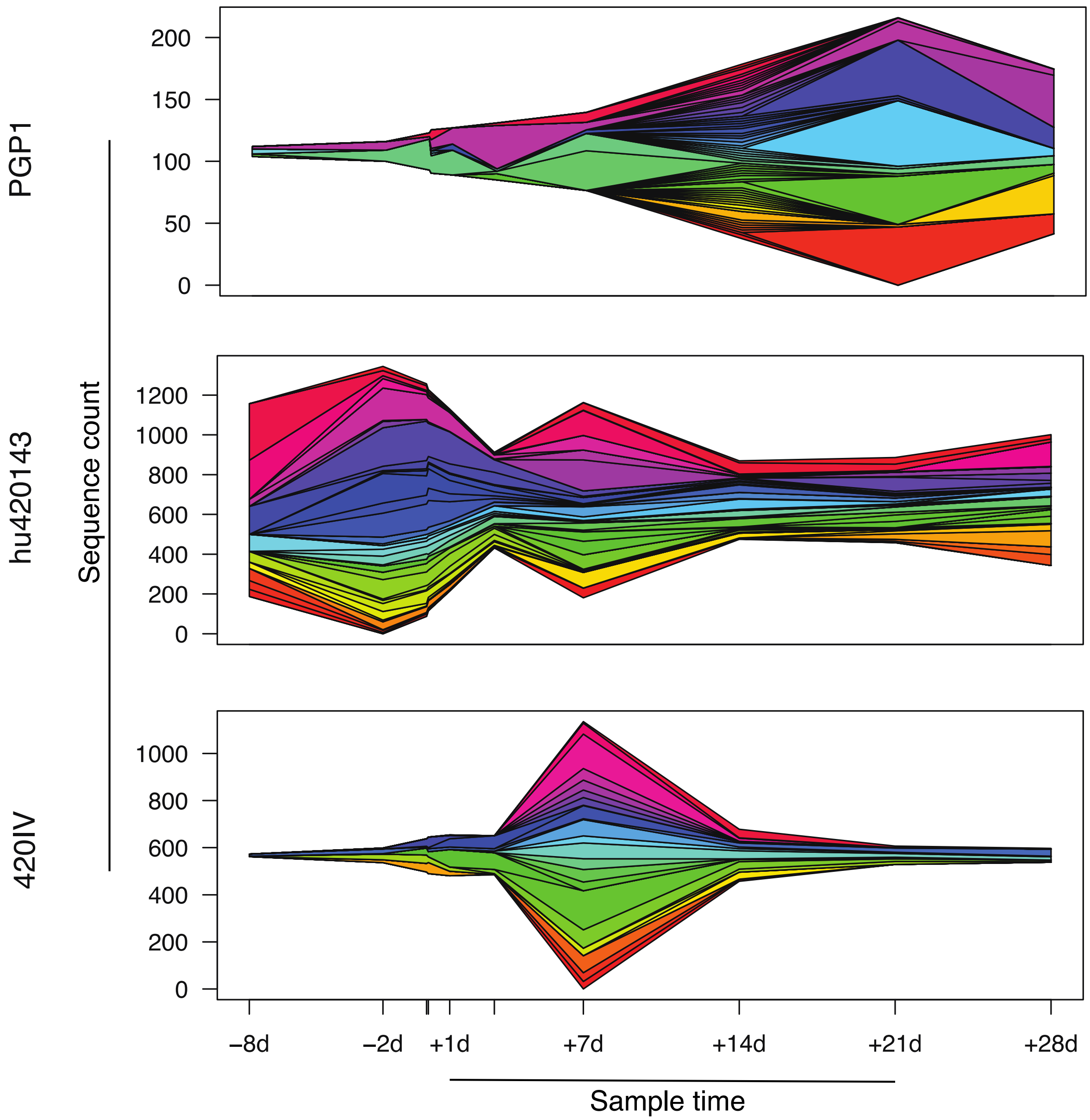


**Supplemental File 8:** Stream plots showing sequence count of large (≥1% of total repertoire sequences) clonal lineages over the course of influenza vaccination. Note that here the definition of sequence uniqueness includes the full V, D, and J segment of the BCR.

**Supplemental File 9: Estimates of $\omega_{CDR}$ on differently sized lineages**

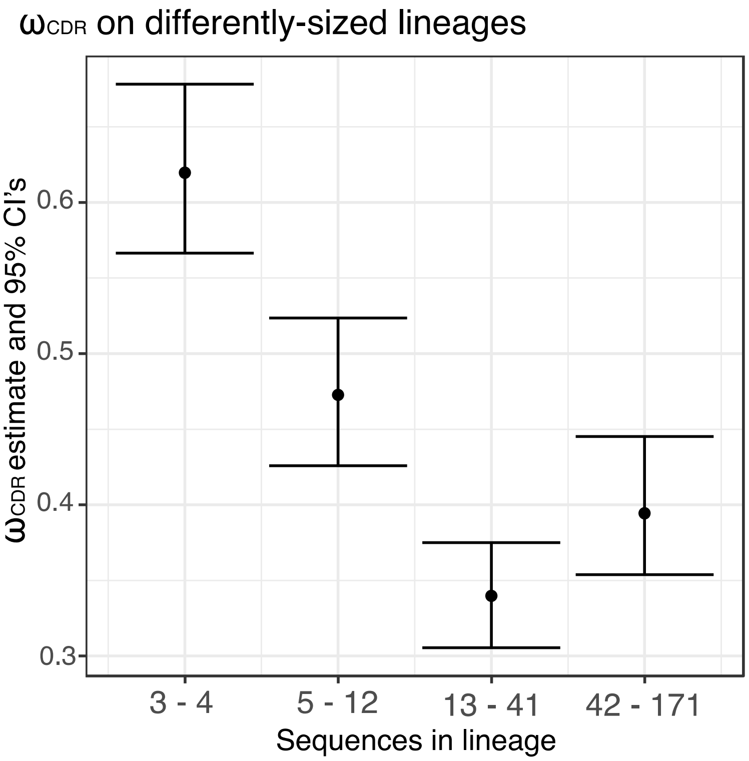


**Supplemental File 9:** Estimates of $\omega_{CDR}$ on differently sized lineages in the same repertoire (subject 420IV, day+7). Samples on the x-axis show approximate quartiles of sequence count in the repertoire, with larger lineages on the right and smaller lineages on the left. Lineages within their respective size constraints were grouped together and $\omega_{CDR}$ was estimated for each group using the HLP19 model and the same parameter estimation procedure described in **Methods***: Phylogenetic model parameter and topology estimation*. These results show that lineages with more sequences tended to have lower $\omega_{CDR}$ estimates than lineages with fewer sequences, though this trend is not monotonic. Sequence counts represent number of unique sequences in a lineage before CDR3 region is masked.
